# Self-reactive B cells traverse a perfect storm of somatic mutagenesis to cause a virus-induced autoimmune disease

**DOI:** 10.1101/2024.01.07.574561

**Authors:** Clara Young, Mandeep Singh, Katherine JL Jackson, Matt A Field, Timothy J Peters, Stefano Angioletti-Uberti, Daan Frenkel, Shyamsundar Ravishankar, Money Gupta, Jing J Wang, David Agapiou, Megan L Faulks, Ghamdan Al-Eryani, Fabio Luciani, Tom P Gordon, Joanne H Reed, Mark Danta, Andrew Carr, Anthony D Kelleher, Gregory J Dore, Gail Matthews, Robert Brink, Rowena A Bull, Daniel Suan, Christopher C Goodnow

## Abstract

The unexplained association between infection and autoimmune disease is strongest for hepatitis C virus-induced cryoglobulinemic vasculitis (HCV-CV). We traced the evolution of the pathogenic rheumatoid factor (RhF) autoantibodies in four HCV-CV patients by deep single cell multi-omic analysis, revealing three sources of B cell somatic mutation converged to drive accumulation of a large disease causing clone. A sensitive method for quantifying low affinity binding revealed three recurring heavy/light chain combinations created by *V(D)J* recombination bound self IgG but not viral E2 antigen. Whole genome sequencing revealed accumulation of thousands of somatic mutations, at levels comparable to CLL and normal memory B cells, but with 1-2 corresponding to driver mutations found recurrently in B cell leukemia/lymphoma. *V(D)J* hypermutation created autoantibodies with compromised solubility. In this virus-induced autoimmune disease, infection promotes a perfect storm of somatic mutagenesis in the descendants of a single B cell.

## Introduction

Viral infection is postulated to be a trigger for many autoimmune diseases, yet their connection is poorly understood. Here, we explore the pathogenesis of one of the strongest known associations between an autoimmune disease and an infection, namely the development of cryoglobulinemic vasculitis (CV) following infection with hepatitis C virus (HCV).

An estimated 58 million people are living with HCV infection^1^, and the development of direct-acting antiviral (DAA) therapy has been a pivotal impetus for the global hepatitis C elimination strategy^2^. HCV-CV is an autoimmune disease complicating HCV infection, characterised by small vessel leukocytoclastic vasculitis, and mediated by a rheumatoid factor (RhF) cryoglobulin: an IgM autoantibody that binds multiple monomers of plasma IgG, forming insoluble precipitates at temperatures below 37°C that redissolve upon warming^3–6^. Strikingly, approximately half of individuals infected with HCV develop a detectable cryoglobulin in the blood. A smaller subgroup develop clinical vasculitis: purpuric skin rashes and arthralgias develop in >15% of HCV-infected individuals, while neuropathy or glomerulonephritis occur in 9% and 5% of cases, respectively^7^. In individuals diagnosed with CV prior to the advent of HCV treatments, >80% had HCV infection and were viremic with HCV RNA^8–10^. In ∼30% of individuals with HCV-CV, the IgM autoantibody is polyclonal (type 3 cryoglobulinemia), while in ∼70% of cases the IgM is monoclonal (type 2 cryoglobulinemia) and associated with higher cryoglobulin levels^7,11^. Individuals with acute HCV-CV are often treated with immunosuppression, but clearance of the virus through DAA treatment results in gradual remission of vasculitis in 70-90% of individuals^12–16^.

Antigenic mimicry between viral antigens and self-antigens is one potential explanation for the association between infection and autoimmunity. In support of this possibility, analysis of large numbers of individual HCV-neutralising antibodies and of HCV-induced cryoglobulin RhFs has revealed recurring sequence mimicry. A single germline antibody heavy chain variable element, *IGHV1-69*, is used by 90% of potent HCV neutralising antibodies, which recognise the HCV envelope glycoprotein, E2^17–23^. The light chains in these antibodies employ a range of kappa light chain variable elements, with 22% employing *IGKV3-20*. Identical pairing of *IGHV1-69* with *IGKV3-20* occurs in 60% of pathogenic IgM RhFs in HCV-CV, comprising the public “Wa” antibody idiotype^24–26^. The fact that virus neutralising antibodies and pathogenic autoantibodies in HCV infection often start out with identical sequences at four of six complementarity determining regions (CDRs) in the antigen binding site supports the antigenic mimicry hypothesis. Expanded B cell clones displaying monoclonal *IGVH1-69/IGKV3-20* IgM receptors on their cell membrane are found in the blood of many individuals with HCV-CV as a non-leukemic, monoclonal or oligoclonal B lymphocytosis^27–32^. These clonal B cell expansions have usually acquired V(D)J mutations required for binding to self IgG as measured by ELISA, but an unresolved question is whether or not the unmutated precursor antibodies bind HCV E2 or IgG^31–35^.

For reasons yet to be determined, individuals infected with HCV are 2.4 times more likely to develop B cell lymphoma than the general population and this increases to 35 times in those with CV^36,37^. While lymphomas are still an uncommon sequelae, those that develop in HCV infected individuals often display membrane IgM with the same public Wa idiotype and RhF activity as the pathogenic secreted cryoglobulins^21,38,39^. One hypothesis is the clones making the pathogenic autoantibody have escaped immune tolerance checkpoints and accumulated to larger numbers by acquiring somatic mutations in B cell regulatory genes, corresponding to recurrent “driver” mutations found in B cell lymphomas and leukemias^40–42^. B cell tolerance checkpoints inhibit self-reactive B cell survival, proliferation and plasma cell differentiation ^42,43^. Lymphoma driver mutations often dysregulate these processes^44^. While it is not known if the cells making pathogenic autoantibody in HCV-CV carry lymphoma driver mutations, such mutations have been found in patients developing CV as a complication of Sjogren’s syndrome^45^, where there is no known infectious trigger.

To investigate how HCV infection may trigger autoimmune CV, here we trace the steps in the evolution of the pathogenic RhF autoantibodies by performing in-depth single cell RNA, DNA and protein analysis of the self-reactive B cell clones in four HCV-CV patients. The results reveal that HCV-CV is caused by a convergence of three sources of somatic mutagenesis.

## Results

### Large IgM^+^ memory B cell clones with public autoantibody idiotypes before and after HCV treatment

In four patients aged 54-66 years diagnosed with HCV-CV involving their skin and nerves, joints or kidneys, peripheral blood was analysed at baseline when HCV RNA was detectable (HCV^+^), and again 25-50 weeks later following DAA therapy-induced viral clearance (HCV^−^) (Figure 1A, Table S1). Following HCV clearance, all four experienced remission of vasculitis symptoms 1-26 weeks after DAA commencement. Patient P1 experienced a relapse of HCV-CV followed by remission again at 124 weeks following DAA commencement (Table S1).

**Figure 1.**
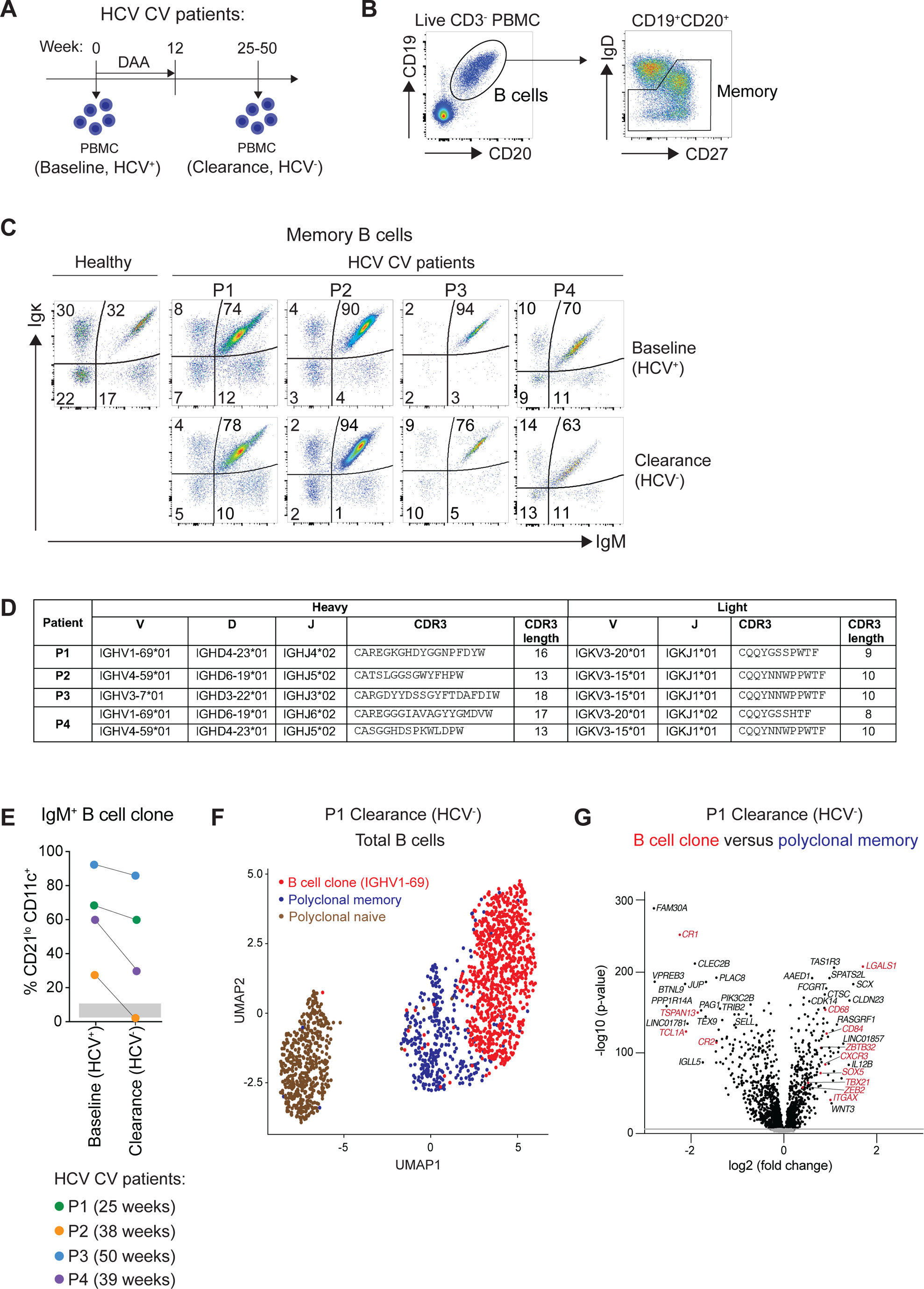
IgM^+^ B cell clonal expansions persist in the blood of cryoglobulinemic vasculitis patients following clearance of HCV. A. Timeline of treatment and sample acquisition. B. Gating strategy for memory B cells. C. Percentage of memory B cells expressing membrane IgM heavy chain and kappa (Igκ) light chains in each patient (P1 to P4) before (HCV^+^) and after virus clearance (HCV^−^). D. Immunoglobulin heavy chain *VDJ* and light chain *VJ* gene rearrangements in each patient’s clone determined by pooled and single cell immunoglobulin RNA sequencing. E. Frequency of CD21^lo^ CD11c^+^ cells within the B cell clone of each patient at baseline and clearance. Grey shading: range of CD21^lo^CD11c^+^ cells amongst memory B cells of three age matched healthy controls. Gating strategy shown in Figure S1F. Number of weeks between baseline and clearance sample as indicated. F. Single cell mRNA sequencing analysis of immunoglobulin receptor and global gene expression in CD19^+^CD20^+^ B cells sorted from the blood of patient P1 at the clearance (HCV^−^) timepoint, with dimension reduction by Uniform Manifold Approximation and Projection (UMAP). Individual cells (denoted by dots) were annotated as the IGHV1-69 B cell clone (*n* = 849, red) or polyclonal cells based on their immunoglobulin sequence. Polyclonal naïve (*n* = 490, brown) and memory B cells (*n* = 360, dark blue) were annotated according to landmark genes. G. Volcano plot of differentially expressed mRNAs between single cells corresponding to the clonal IGHV1-69 B cells (*n*= 849) and the polyclonal memory B cells (*n*=360) of patient P1 at the clearance timepoint. Differentially expressed genes with family-wise error rate (FWER) <0.05 are shown in black. Landmark mRNAs increased (e.g. *ITGAX*, *TBX21*) or decreased (e.g. *CR2*) in CD21^lo^ CD11c^+^ age-associated memory B cell genes are shown in red. Immunoglobulin V genes were excluded from the analysis.

Circulating B cell clones were identified in each patient both before and after HCV clearance, defined as CD19^+^ CD20^+^ IgD^low/-^ memory B cells (Figure 1B) expressing surface IgM and restricted to kappa light chain (IgM^+^ IgK^+^; Figure 1C). At the baseline timepoint when the patients had active HCV infection (HCV^+^), the IgM^+^ IgK^+^ B cell clone comprised between 70-94% of the memory B cell compartment (Figure 1C, top panel), which remained largely unchanged in all four patients following clearance of HCV (HCV^−^) (Figure 1C, bottom panel). No patients exhibited an elevated total lymphocyte count (Table S1) or an overall increased frequency of B cells, with the exception of patient P2 who demonstrated a slightly elevated B cell frequency at the clearance (HCV^−^) timepoint (Figure S1A).

Bulk and single cell immunoglobulin RNA-sequencing of the patients’ blood confirmed the expanded IgM^+^ B cell clones at the clearance timepoint (HCV^−^) were the same B cell clones detected at the baseline timepoint (HCV^+^) for each patient (Figure S1C,D). The IgM^+^ B cell clones of patients P1, P2 and P3 were monoclonal and corresponded to recurring “public” idiotypes of HCV-associated RhF immunoglobulin heavy and light chain combinations encoded by *IGHV1-69* and *IGKV3-20* (Wa idiotype), *IGHV4-59* and *IGKV3-15* (Bla idiotype), or *IGHV3-7* and *IGKV3-15* (Po idiotype) (Figure 1D). The IgM^+^ B cell expansion of patient P4 was bi-clonal, composed of a dominant B cell clone with *IGHV1-69/IGKV3-20* immunoglobulin comprising 35% of total B cells at baseline and a smaller second B cell clone with a *IGHV4-59/IGKV3-15* immunoglobulin comprising 3% of total B cells at baseline (Figures 1D, S1C, S1D, data not shown).

Using the anti-idiotypic monoclonal antibody, G6, specific for immunoglobulins employing the *IGHV1-69* F-type allele^46^, the frequency of the IgM^+^ *IGHV1-69* B cell clone in the blood of patient P4 decreased by 30% 39 weeks after the commencement of DAA therapy, whereas the B cell clone of patient P1 increased by 20% 13 weeks post DAA therapy (Figure S1B). While it is difficult to determine the precise shifts in B cell clone frequencies between the two blood sample timepoints, the expanded B cell clones persisted in the blood up to a year following DAA commencement despite sustained clearance of the virus.

### Persistence of CD2l^lo^ CD11c^+^ age-associated memory B cell clones in the absence of virus

At the baseline HCV^+^ timepoint, the majority of cells comprising the IgM^+^ B cell clones in patients P1, P3 and P4 (and 28% of the clone in P2) displayed a CD21^lo^\CD11c^+^ CD19^hi^ age-associated memory B cell profile (Figures 1E, S1E and S1F), consistent with previous studies of HCV-CV patients^28,31,32,34^. This subset is considered to be a population of memory B cells poised for plasma cell differentiation and antibody secretion^47^, and normally comprises less than 10% of memory B cells in healthy donors (Figure 1E). At the clearance timepoint, there was a small decrease in frequency of CD21^lo^ CD11c^+^ cells in patients P1, P3 and P4, which continued to constitute 59%, 87% and 35% of the clone, respectively (Figure 1E). Single cell Repertoire And Gene Expression by mRNA sequencing (RAGE-seq)^48^ analysis of B cells from patient P1 at the clearance (HCV^−^) timepoint confirmed the CD21^lo^ CD11c^+^ memory B cell profile in the persisting IGHV1-69^+^ IgM^+^ B cell clone with upregulation of landmark genes including *ITGAX* (CD11c), *LGALS1* (Galectin-1), *CXCR3*, *SOX5*, *ZBTB32*, *TBX21* (T-bet) and *ZEB2* (Figure 1H, Table S2), indicating the virus is not required for sustaining the age-associated memory B cell phenotype of the expanded clones. Patient P2 was the only case where the CD21^lo^ CD11c^+^ subset declined into the normal range, despite the clone persisting or increasing in this patient at the clearance timepoint (Figure 1C).

### The landscape of somatic mutations across the genome of HCV-CV clonal expansions

To identify genome-wide somatic mutations that might confer a clonal advantage, thousands of cells from each patient’s expanded IgM^+^ IgK^+^ B cell clone at the baseline timepoint (HCV^+^) were bulk sorted with estimated clone purity >90% and genomic DNA processed for short-read whole genome sequencing (WGS) (Figure 2A). Paired analysis of WGS on DNA from pools of sorted control B cells from each patient (Figure 2B) was performed to remove germline variants. A three-tiered bioinformatic approach was employed to detect somatic mutations: i) SNVs and small (<50bp) indels; ii) large chromosomal SVs including >50bp deletions, translocations, inversions; and iii) somatic ploidy events (Figure 2A). As an internal benchmark, the same bioinformatic pipeline called somatic mutations in tumour/normal WGS pairs from four *IGHV*-mutated B cell CLLs and four B cell non-Hodgkin’s lymphomas (B-NHL) representative from a large published analysis^49^.

**Figure 2.**
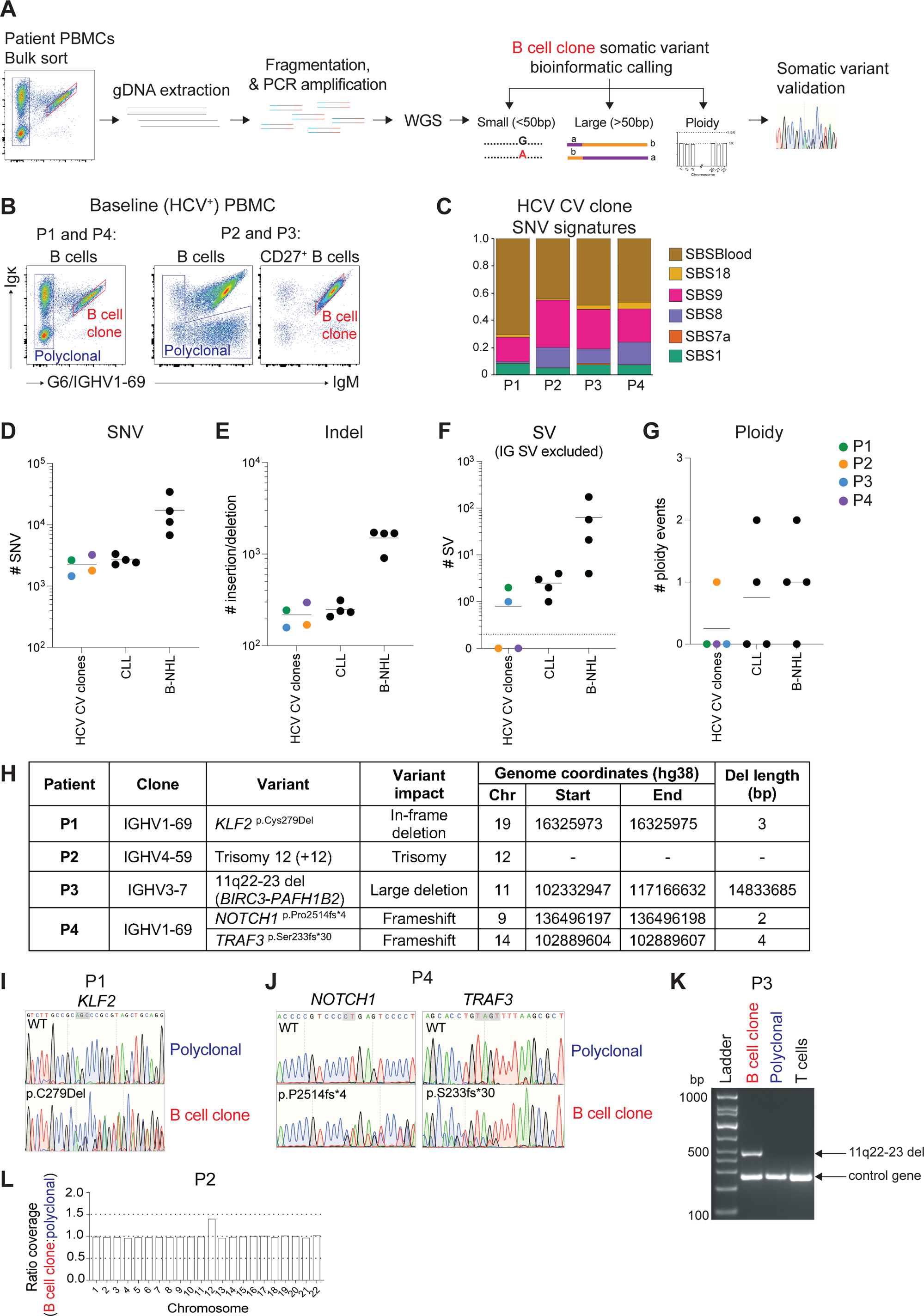
Somatic genome-wide and lymphoma driver mutations in the IgM^+^ clone of each patient. A. Workflow to detect genome-wide somatic mutations. B. Gating strategy for bulk sorting of clonal B cells and polyclonal memory B cells from patient blood at the baseline (HCV^+^) timepoint. C. Proportion of somatic single nucleotide variants (SNV) in each HCV-CV patient’s clone corresponding to the indicated mutational signatures. D-G. Number of somatic single nucleotide variants (SNV) (D), indels (E) structural variants (SV) at non-immunoglobulin (IG) loci (F) and ploidy events including chromosome gain/loss (G) within the whole genome sequence of the HCV-CV B cell clones (P1-P4) compared to 4 representative IGHV-mutated CLL samples and 4 B cell non-Hodgkin’s lymphomas^49^. Dotted line in (F) indicates change from logarithmic scale to linear scale on the y-axis. H. Summary of lymphoma driver mutations detected. I. Confirmation of *KLF2^p.C279del^* somatic mutation in patient P1’s clone and absence in polyclonal B cells by PCR and Sanger sequencing. J. Confirmation of the *NOTCH1^p.P2514fs*^*^4^ and *TRAF3^p.S233fs*^*^30^ somatic mutations in patient P4’s clone by PCR and Sanger sequencing. K. Confirmation of the 11q22-23 deletion in the clone (red) and not in polyclonal B cells (blue) or T cells of patient P3 by PCR amplification across the breakpoint (top arrow). Primers to a control gene (*SEC23IP*) were included as a positive control (bottom arrow). L. Ratio of average WGS read depth per chromosome in patient P2’s clone relative to polyclonal memory B cells.

Across the genome, the HCV-CV B cell clones carried a mean of 2281 SNVs (range 1459 – 6771) and 217 indels (range 169-296; Figures 2D, E, Table S3). A comparable mutation burden of mean 2678 SNVs and 248 indels was called in the four CLLs, consistent with independent analysis of a large set of CLLs^49^, whereas SNV and indel burdens were 7-8 times higher in B-NHLs^49^. The majority of SNVs in the HCV-CV clonal B cells corresponded to mutation signatures associated with blood stem cell ageing (SBSblood) and non-canonical activation-induced cytidine deaminase activity (SBS9) (Figure 2C), as observed in *IGHV*-mutated CLL^49,50^ and in normal memory B cells^51,52^. The HCV-CV B cell clones carried a mean of 0.75 (range 0-2) chromosomal structural variants not involving immunoglobulin genes (Figure 2F, Table S4), slightly lower than the burden in CLL (mean 2.5, range 1-4) and much lower than in B-NHL (mean 64, range 4-174) (Figure 2F). With respect to ploidy events, only the HCV-CV P2 clone had an event detected consisting of a whole chromosome 12 gain (Figure 2G, Table S5). A similar burden of ploidy events was called in CLL and B-NHL (range 0-2), including the identical chromosome 12 gain in two of the CLL samples (Table S5). Collectively these analyses suggest the overall somatic mutation burden of the HCV-CV clones in these four patients is similar to CLL and significantly lower than B-NHL.

### Lymphoma driver somatic mutations in each patient’s B cell clone

Cross-referencing the list of somatic mutations in each HCV-CV B cell clone with databases of lymphoma driver gene somatic mutations found recurrently in human B cell lymphomas and leukemias (Table S6) revealed a single putative driver mutation in patients P1, P2 and P3 and two putative driver mutations in P4 (Figure 2H). In patient P1, a heterozygous 3bp exonic deletion present in 35% of reads, *KLF2*^p.Cys279Del^, deleted the cysteine at position 279 (Figures 2I, S2B, data not shown). Loss-of-function *KLF2* somatic mutations occur in 20-42% of splenic marginal zone lymphoma (SMZL) and ∼30% of diffuse large B cell lymphomas (DLBCL) associated with hepatitis B virus infection (Figure S2B)^53–56^. The deleted cysteine is absolutely conserved between species and paralogues (Figure S2C) and critical for coordinating a zinc ion to fold the C2H2 zinc finger for DNA binding. A *KLF2*^p.Cys274Tyr^ somatic mutation in the other zinc-coordinating cysteine in this finger, from a SMZL patient, has been shown to confer loss-of-function^54^. We confirmed both *KLF2*^p.Cys279Del^ and *KLF2*^p.Cys274Tyr^ mutations diminish the ability of KLF2 to repress an NF-κB reporter in transfected cells (Figure S2D).

In patient P4, the clone had two driver somatic mutations: a 2 nucleotide exonic deletion, *NOTCH1*^p.Pro2514fs*4^, and a 4 nucleotide exonic deletion, *TRAF3*^p.Ser233fs*30^, present in 45% and 39% of reads, respectively (Figures 2J, S2E, S2F, data not shown). *NOTCH1* ^p.Pro2514fs*4^ occurs in 4-12% of CLL patients and in monoclonal B cell lymphocytosis (MBL), conferring gain-of-function by truncating the C-terminal PEST domain that targets the NOTCH1 intracellular domain for proteasomal degradation (Figure S2E)^57–63^. The *TRAF3*^p.Ser233fs*30^ mutation truncates the protein, eliminating the coiled-coil domain required for heterodimerization with TRAF2 and stabilising TRAF3 trimers^64^ and eliminating the MATH/TRAF-C domain that recruits NIK for ubiquitination by BIRC3^65,66^. Similar *TRAF3* truncating mutations and small chromosomal deletions that eliminate one copy of *TRAF3* occur in 10% of SMZL, 2% of CLL, 10% of myelomas, 9% of DLBCL, and ∼45% of canine B cell lymphomas^59,67–71^.

In patient P3’s clone, *BIRC3* was inactivated by a 15 Mb deletion within chromosome 11 (11q22-23 del) with breakpoints in *BIRC3* and *PAFH1B2* (Figure 2K, S2I Table S4). The presence of the deletion in clonal B cells but not polyclonal memory B cells or T cells was confirmed by PCR with primers flanking the breakpoint (Figure 2K). Between 18 and 30% of CLL cases carry 11q deletions centred on loss of *ATM*, encoding a master regulator of the DNA damage response pathway^59,72^. Patient P3’s 11q22-23 deletion results in a truncating mutation, *BIRC3*^p.N442*^, and heterozygous loss of 95 protein coding genes including genes relevant to B cell function: *ATM*, *BIRC2*, *POU2AF1*, *BTG4*, *CUL5*, *NPAT*, *PPP2R1B* and the inflammatory caspases (caspases 1, 4 and 5) (Figure S2I). *BIRC3* loss-of-function mutations occur in 11% of SMZL, 5% of CLL and 8% of MBL, and loss of *BIRC3* occurs in 83% of CLL cases with an 11q deletion^57,59,70,73^.

Patient P2’s clone had chromosome 12 trisomy (+12) (Figure 2L and Table S5), a cytogenetic abnormality found in 10-20% of CLL^59,72^ (Table S5) and 18% of people with MBL and normal lymphocyte counts^74^. WGS read coverage was increased 1.4-1.5X for chromosome 12 from sorted clonal cells compared to their polyclonal memory B cell counterparts, both in the baseline blood sample (Figure 2L) and after HCV clearance (Figure S2G, Table S5). GSEA of differentially expressed mRNAs in clonal B cells revealed enrichment of gene sets corresponding to many chromosome 12 cytogenetic bands (Figure S2H), as observed for CLL samples with trisomy 12^75,76^.

### Lymphoma driver mutations arise prior to clonal tree branching

Amplification and sequencing of mRNA and genomic DNA from single sorted memory B cells (Figure 3A) was used to analyse intraclonal heterogeneity of the HCV-CV clonal B cells at the baseline timepoint. The *V(D)J* sequences of heavy and light chain mRNAs revealed prior and ongoing somatic hypermutation amongst the clonal cells, including multiple amino acid substitutions in the CDRs. Stepwise accumulation of individual *V(D)J* mutations in different cells within the clone enabled generation of a clonal tree for each patient (Figures 3B-3E). Paired analysis of amplified genomic DNA for a subset of these single B cells enabled the driver mutations to be placed within the clonal trees. As controls, sorted single memory B cells that were not part of the expanded clone were analysed in parallel, with each displaying a polyclonal, unique antibody *V(D)J* rearrangement (Figure S3).

**Figure 3.**
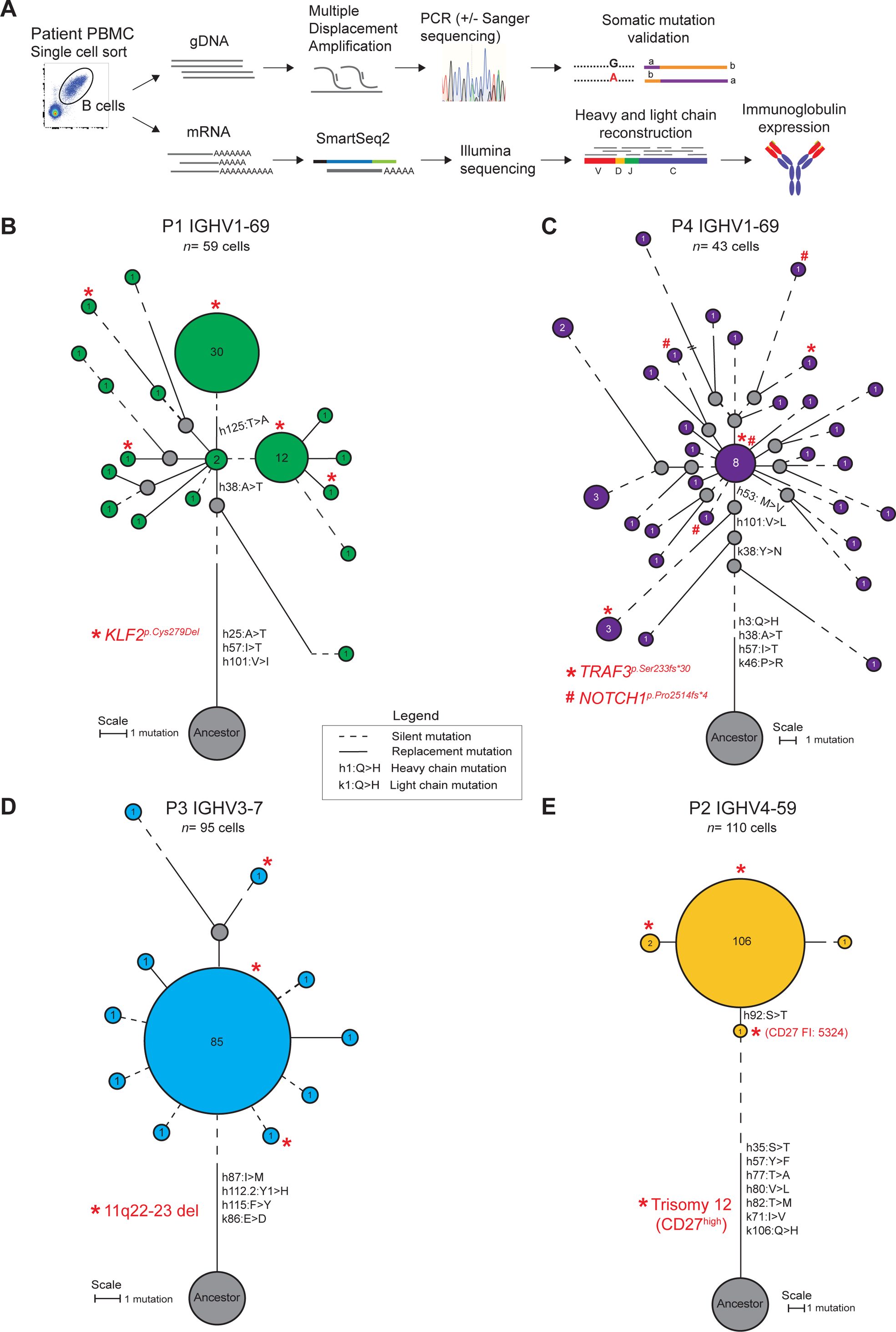
Clonal trees inferred from single cell mutation analysis. A. Schematic of mutation detection in genomic DNA and immunoglobulin *V(D)J* mRNA. B-E. Clonal trees depicting the acquisition of individual *V(D)J* mutations by manual nearest neighbour analysis. Grey circles indicate inferred unmutated ancestor (“ancestor”) and inferred intermediate cells. Coloured circles denote one or more sequenced cells sharing the identical immunoglobulin nucleotide sequence, with size of circle proportional to number of cells (cell number indicated in centre). Red asterix or hash denotes one or more cells identified with the indicated driver mutation. Length of branches corresponds to the number of *V(D)J* mutations (see scale): solid lines, number of replacement mutations; dotted lines, number of silent mutations. Amino acid change and position (IMGT numbering) are indicated with lower case “h” for heavy chain and “k” for the kappa light chain. Trisomy 12 in single cells from patient P2 was inferred from flow cytometric measurement of high CD27.

In patient P1 (Figure 3B), the *KLF2*^p.Cys279Del^ mutation was detected in cells across the clonal tree including branches that precede the predominant expansion of cells with the heavy chain Thr125Ala (h125:T>A) mutation that increases binding to IgG (see below). The mutant *KLF2* allele was amplified in 63% of individual clonal B cells tested (*n*=7/11 cells) and none of the polyclonal memory B cells (*n*=0/16 cells) (Figure S3A). Only ∼50% of allelic sequences present in genomic DNA of single cells are amplified during multiple displacement amplification, explaining why the mutant allele was not detected in all cells from the clonal tree. Thus, the *KLF2* mutation was acquired before the clone acquired potent RhF activity and accumulated to the size circulating when P1 began DAA treatment.

Similarly, in patient P4, the *NOTCH1*^p.Pro2514fs*4^ and *TRAF3*^p.Ser233fs*30^ mutations were both found in single cells across the clonal tree, but not in polyclonal memory B cells (Figure 3C, Figure S3A). *TRAF3*^p.Ser233fs*30^ was already present in an early branch preceding acquisition of the heavy chain M53V mutation shared by 88% of clonal B cells (Figure 3C).

By contrast, the pathogenic clones in patients P2 and P3 (Figures 3D and 3E) displayed almost no intraclonal *V(D)J* diversity. PCR amplification of single cells from patient P3 with flanking primers detected the 11q22-23 deletion in 60% of individual clonal B cells (*n*=18/30 cells) and none of three polyclonal B cells sampled (*n*=0/3 cells) (Figure S3A) consistent with carriage by all of the clonal cells. In patient P2, flow cytometric measurement of CD27 (encoded on chromosome 12) revealed increased CD27 on the majority of clonal cells compared to polyclonal memory B cells (Figure S3B). CD27 was not increased on clonal cells of the other patients (data not shown). In P2, CD27 fluorescence intensity >2 s.d. above the mean of non-expanded polyclonal B cells was observed on 55% of clonal cells (n=33/60 cells) including an ancestral cell yet to acquire the heavy chain Ser92Thr (h92:S.T) mutation shared by all other cells in the circulating clone (Figures 3D and S3B). Collectively, these results indicate that a lymphoma driver gene mutation was acquired before each clone accumulated to the large size circulating at the baseline timepoint.

### Accumulation of *V(D)J* mutations confers pathogenic autoantibody activity

Next, we tested the hypothesis that the IGHV1-69^+^ IgK^+^ B cells (P1 and P4) and IgM^+^ IgK^+^ B cells (P2 and P3) were the source of RhF cryoglobulins in the HCV-CV patients. We selected the immunoglobulin sequence shared by at least 50% of individual clonal cells; Figures 3B-3E, S4A and S4B) and expressed it as pentameric secretory IgM with J-chain to test for both cryoglobulin and RhF activity. In parallel, unmutated ancestor IgM antibodies were synthesised corresponding to each patient’s most frequent clone but with the V(D)J somatic mutations reverted to their ancestral sequence (Figure S4A and S4B).

The predominant mutated clonal IgM antibodies from patients P1, P2, and P4 bound IgG in ELISA and thus demonstrated RhF activity (Figure 4A), and demonstrated cryoglobulin activity by forming precipitates with IgG at temperatures below 25°C (Figure 4B). By contrast, IgM corresponding to the unmutated ancestor of each clone displayed minimal or no detectable RhF or cryoglobulin activity (Figures 4A and 4B). These results indicate the 5-8 non-synonymous V(D)J mutations acquired by the IgM^+^ B cell clones in patients P1, P2 and P4 conferred both self-reactivity against IgG and pathogenic insolubility of the IgM autoantibody.

**Figure 4.**
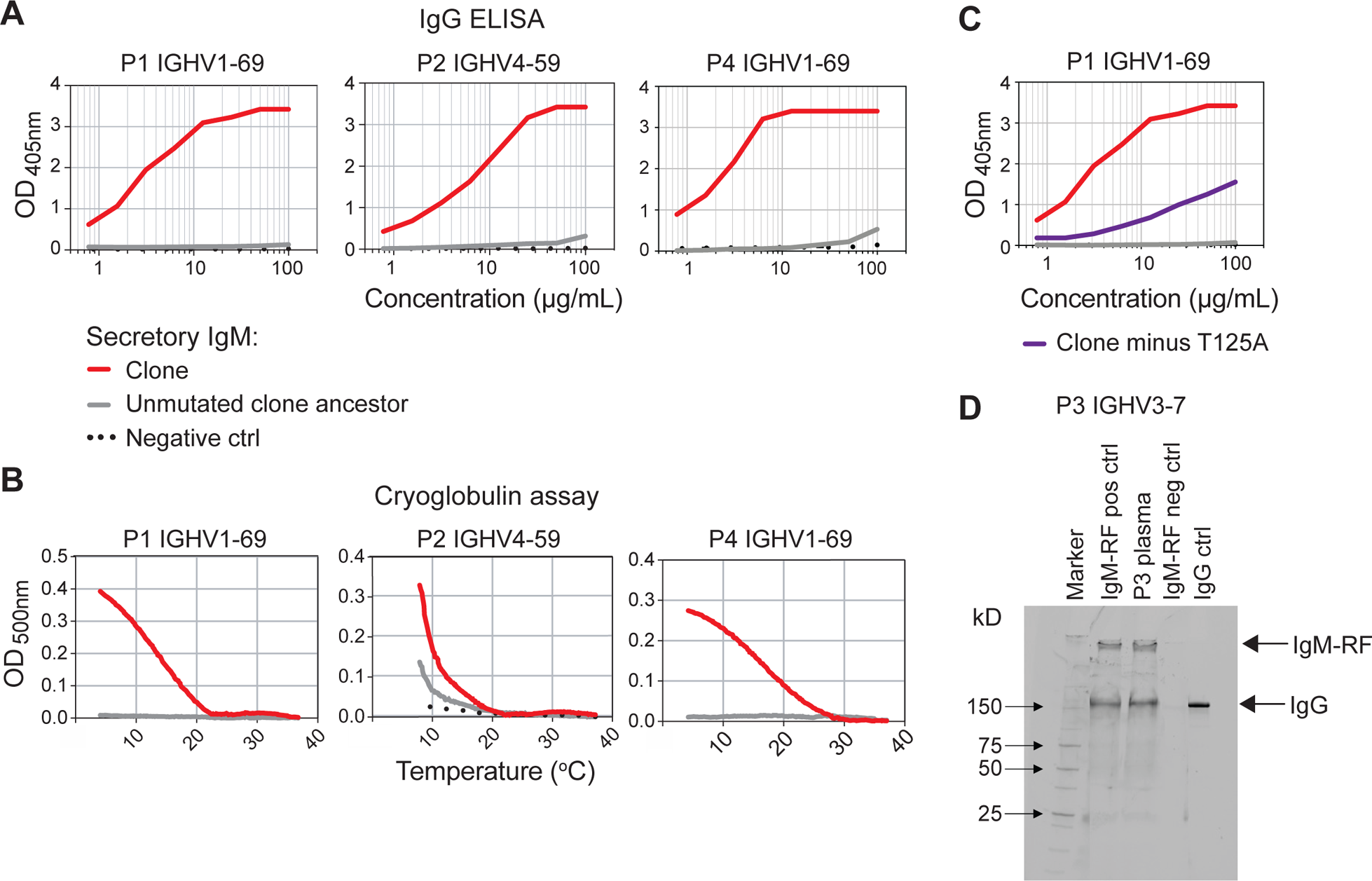
IgG binding and cryoprecipitation of expressed pentameric IgM. A. Pentameric secreted IgM corresponding to the predominant clonal sequence (red) and unmutated ancestor (grey) of each patient was expressed *in vitro* and tested by ELISA at the indicated concentrations for binding to immobilised human IgG. Negative control (black) was anti-HCV broadly neutralising antibody AR3C expressed as pentameric secreted IgM. Representative of 2-4 independent experiments, run in duplicate. B. Pentameric secreted IgM antibodies expressed as in (A) were mixed with human polyclonal IgG and aggregation measured by visible wavelength optical density in a temperature-controlled spectrometer cooled from 37°C to 4°C at 0.1°C/min. Representative of 2-3 independent experiments. C. ELISA for expressed IgM from patient P1 as described in A, but comparing IgM corresponding to the predominant clonal sequence without the IGHV1-69 T125A mutation (dark purple). D. Heat-aggregated IgG was mixed with plasma from patient P3 at baseline timepoint, or positive or negative control plasma samples, and precipitates analysed by SDS-PAGE under non-reducing conditions to detect intact IgM and IgG (arrows).

In patient P1, 51% of the clonal cells had diverged from their siblings by acquiring a heavy chain Thr125Ala mutation (Figure 3B). When this single mutation was reverted to express the immediate precursor antibody, corresponding to that made by 24% of clonal cells (*n*=14/59 cells), the RhF activity measured by ELISA was markedly reduced (Figure 4C, S4C). Since cells in both branches already carried *KLF2*^p.Cys279Del^, the main population of clonal cells in patient P1 acquired increased self-reactivity to IgG after acquisition of the somatic driver mutation.

Two heavy chain mutations, Ala38Thr and Ile57Thr, were independently acquired early in the clonal tree by the *IGHV1-69* B cell clones of patient P1 and P4 (Figure S4A). Alternate versions of each patient’s IgM antibody were expressed individually reverting the Ala38Thr or Ile57Thr mutations while maintaining all other mutations, with the exception of the Thr125Ala mutation in patient P1 (discussed above), which was also reverted. Reversion of the Ala38Thr mutation had no significant impact on IgG binding (Figure S4C). By contrast, reversion of the Ile57Thr mutation abolished IgG binding as measured by ELISA for the clonal antibodies of patients P1 and P4 (Figure S4C), consistent with convergent evolution to increase self-reactivity. It is notable that the Ile57Thr (h57:I.T) mutation arose early in the clonal tree in both patients (Figures 3B and 3C).

In contrast to the other three patients, the synthesised IGHV3-7 IgM of patient P3 failed to demonstrate detectable RhF in ELISA (Figure S4D) nor cryoglobulin activity in the rapid cooling spectrophotometer assay (data not shown). However, when the plasma of patient P3 was mixed with heat-aggregated IgG, IgM^+^ RhF activity was detected (Figure 4D). This precipitating IgM^+^ RhF was gel purified and subject to *de novo* sequencing by mass spectrometry to identify clonotypic CDR3 peptides (Figure S5). The heavy and light chain CDR3 amino acid sequences determined by mass spectrometry were identical to the CDR3 sequences of the synthesised IgM, indicating the IGHV3-7/IGKV3-15 antibody expressed on the IgM^+^ B cell clone of patient P3 corresponds to the cryoglobulin autoantibody present in blood. As described below, the IGHV3-7 IgM from patient P3 appears to have lower affinity for IgG than the IgM’s from the other three patients, which may explain the absence of RhF activity by ELISA.

### HCV E2 reactivity in anti-E2 ancestors but not autoantibody ancestors

The results above, showing apparent lack of substantial self IgG binding by the unmutated antibody corresponding to the ancestor B cell for each of the four patients, posed a dilemma for understanding how clonal expansion and somatic *V(D)J* mutation was initiated. Given shared IGHV1-69/IGKV3-20 use by many HCV neutralising antibodies, we sought to distinguish between two possible explanations: (1) the ancestor IgM bound HCV E2 and this initiated B cell proliferation, with daughter cells subsequently acquiring IgG binding through *V(D)J* mutations; or (2) the ancestor IgM did not bind HCV E2 but rather bound IgG, albeit with too low affinity for detection by ELISA. To resolve between the two alternatives, we built upon a protocol that permits detection of low affinity IgM binding to influenza haemagglutinin^77^, by transiently expressing the unmutated ancestor antibodies as membrane IgM on the surface of HEK293 cells (Figure 5A). Binding to multimerised HCV E2 or IgG antigen was tested by flow cytometry using fluorescent dextran molecules carrying ∼25 biotin acceptor sites loaded with biotinylated antigen (Figure 5A). We discovered that a valuable characteristic of this method arises from each individual transfected cell expressing a different amount of membrane IgM on their cell surface, spanning a 500-fold range comparable to the range of membrane IgM concentrations on human B cells (Figure 5B and S6A).

**Figure 5.**
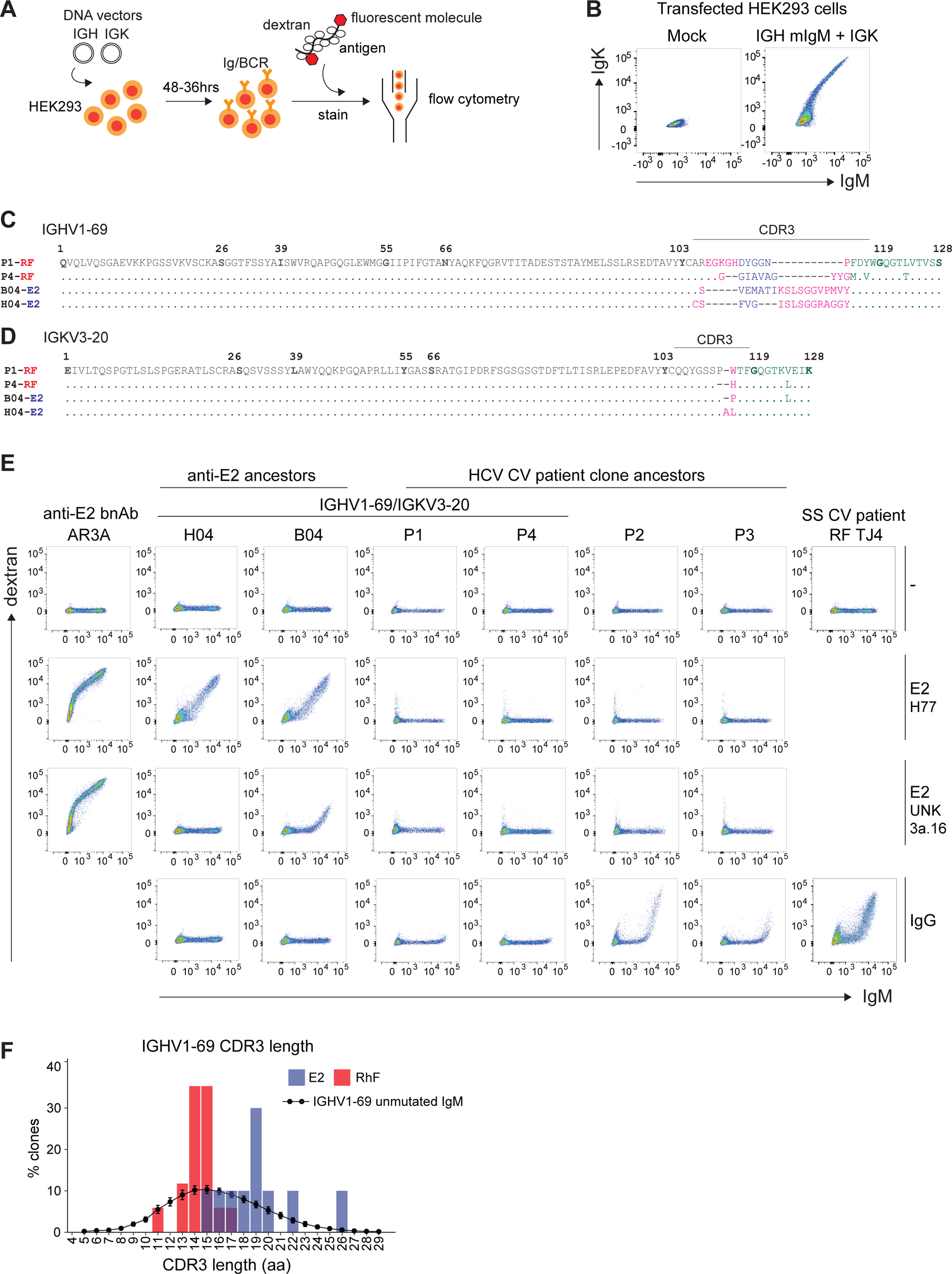
Cryoglobulin clone ancestral IgM bind multimerised IgG but not HCV E2. A. Experimental protocol. HEK293F cells were transiently transfected with vectors encoding membrane IgM corresponding to patient or control antibodies. Fluorescent dextran molecules bearing ∼25 biotin acceptor sites were mixed with biotinylated proteins (e.g. IgG), and binding of the resulting multimeric antigen to the transfected cells analysed by flow cytometry. B. Flow cytometric analysis of transfected or mock-transfected HEK293F cells stained with fluorescently labelled anti-human IgM and anti-human kappa light chain antibodies, demonstrating the wide range of cell surface IgM density from cell to cell. C. Binding of empty dextran (top row), HCV E2 H77-dextran (second row) and HCV E2 UNK3a.13.6-dextran E2-dextran (third row) and IgG-dextran (bottom row) to transfected HEK293 cells expressing membrane IgM corresponding to the mutated anti-HCV E2 broadly neutralising antibody AR3A (far left column), the unmutated ancestors of two anti-HCV E2 antibodies (B04 and H04), the unmutated ancestors of patient P1-P4, or a mutated IgM RhF cryoglobulin from a Sjogrens’s syndrome patient with vasculitis (SS CV) (patient TJ) (far right). D. and E. Amino acid alignments of heavy (D) and kappa light chain (E) variable domains of the unmutated ancestor IgM in patients P1 and P4 and the unmutated ancestor IgM for the memory B cell anti-HCV E2 antibodies B04 and H04. IMGT numbering, with CDR flanking residues in bold. The V gene segment is shown in dark grey, the D gene segment is shown in navy blue, the J gene segment is shown in green and likely N-additions are shown in pink. Periods (.) indicates identity and dashes (-) indicate gaps. F. Percent of IGHV1-69/IGKV3-20 antibodies with the indicated amino acid length of heavy chain CDR3, for antibodies that bind self IgG (RhF, red, *n*=17) and antibodies that bind HCV E2 (E2, blue, *n*=10). The black line denotes corresponding length distribution for 1343 IGHV1-69 IgM antibodies of unknown specificity, representing the naïve repertoire (<2% SHM) from PBMCs of 61 healthy donors. Points show mean ± STD dev.

Transfected cells were tested for binding to biotinylated IgG or E2 multimerised on dextran. HCV E2 corresponded to Genotype 1a (isolate H77) and Genotype 3a (isolate UNK 3a.13.6) which collectively comprise >80% of Australian HCV infections^78^. All of the patients were infected with HCV Genotype 1, with patients P2 and P4 matching the genotype subtype of isolate H77 (Table S1). As positive and negative controls, HEK293 cells were transfected in parallel with vectors encoding membrane IgM corresponding to a hypermutated, pathogenic IGHV1-69/IGKV3-20 IgM cryoglobulin, TJ4, from a Sjogren’s syndrome patient with CV^45^, or a hypermutated, broadly neutralising IGHV1-69/IGKV3-20 HCV E2 antibody, AR3A^19^. In parallel, E2-dextran binding was tested on transfected cells expressing the unmutated ancestor IgM of two clonally unrelated IGHV1-69/IGKV3-20 anti-HCV E2 antibodies, B04 and H04, isolated from circulating memory B cells of a patient chronically infected with HCV (Bull et al, manuscript in preparation). The heavy and light chain amino acid sequences of the ancestral B04 and H04 antibodies were identical to the RhF ancestors of patient P1 and P4 with the exception of the heavy and light chain CDR3 loops (Figures 5C, D).

Cells with each different antibody sequence were transfected, stained and analysed in parallel to hold key variables constant, including extent of antigen multimerization or sensitivity of the flow cytometer. None of the transfected cells, even those with the highest amount of membrane IgM, bound fluorescent dextran that had not been decorated with antigen (Figure 5E, top row). As expected, cells expressing the HCV E2 antibody AR3A bound E2-dextran (Figure 5E, far left column). We found cells expressing membrane IgM with the ancestral sequence of B04 and H04 bound fluorescent dextran decorated with E2 H77 (Genotype 1a) (Figure 5E, second row). A higher threshold membrane IgM was needed for ancestral B04-expressing cells to bind E2-dextran for isolate UNK3a.13.6 (Genotype 3a) (Figure 5E, third row). Strikingly, neither isolate of HCV E2-dextran bound to cells expressing the unmutated ancestor IgM for any of the cryoglobulins from patients P1-P4 (Figure 5E).

By contrast, cells expressing the unmutated ancestor IgM cryoglobulins bound IgG-dextran (Figure 5E, bottom row, Figure 6A). Reciprocally, no IgG-dextran binding was detectable on cell expressing the unmutated ancestral anti-E2 B04 and H04 antibodies (Figure 5E, bottom row). The ancestral B04 and H04 antibodies have longer heavy chain CDR3 loops than the ancestral P1 and P4 antibodies (22 and 19 aa compared to 16 and 17 aa, respectively; Figure 5C). We therefore compared heavy chain CDR3 lengths in a larger set of antibodies by assembling published sequences for IGHV1-69/IGKV3-20 hypermutated antibodies known to bind either HCV E2 (10 antibody sequences) or self-IgG (17 antibody sequences), and for 1343 IGHV1-69 unmutated IgM antibodies of unknown specificity expressed in the circulating B cell repertoire of healthy adults (Figure 5F and Figure S6B). The naïve IgM repertoire of IGHV1-69 antibodies spanned a broad range of heavy chain CDR3 lengths with mode of 15 amino acids. By contrast, the IgG-binding autoantibodies were skewed towards the shorter part of this range (mode 14-15 aa, range 11-17 aa), whereas the HCV E2-binding antibodies were skewed towards longer CDR3 lengths (mode 19 aa, range 15-26 aa) (Figure 5F). Heavy chain CDR3 length, which becomes fixed by V(D)J recombination at the pre-B cell stage, appears to be one of the determining factors for self or virus antigen to bind to IGHV1-69/IGKV3-20 antibodies.

**Figure 6.**
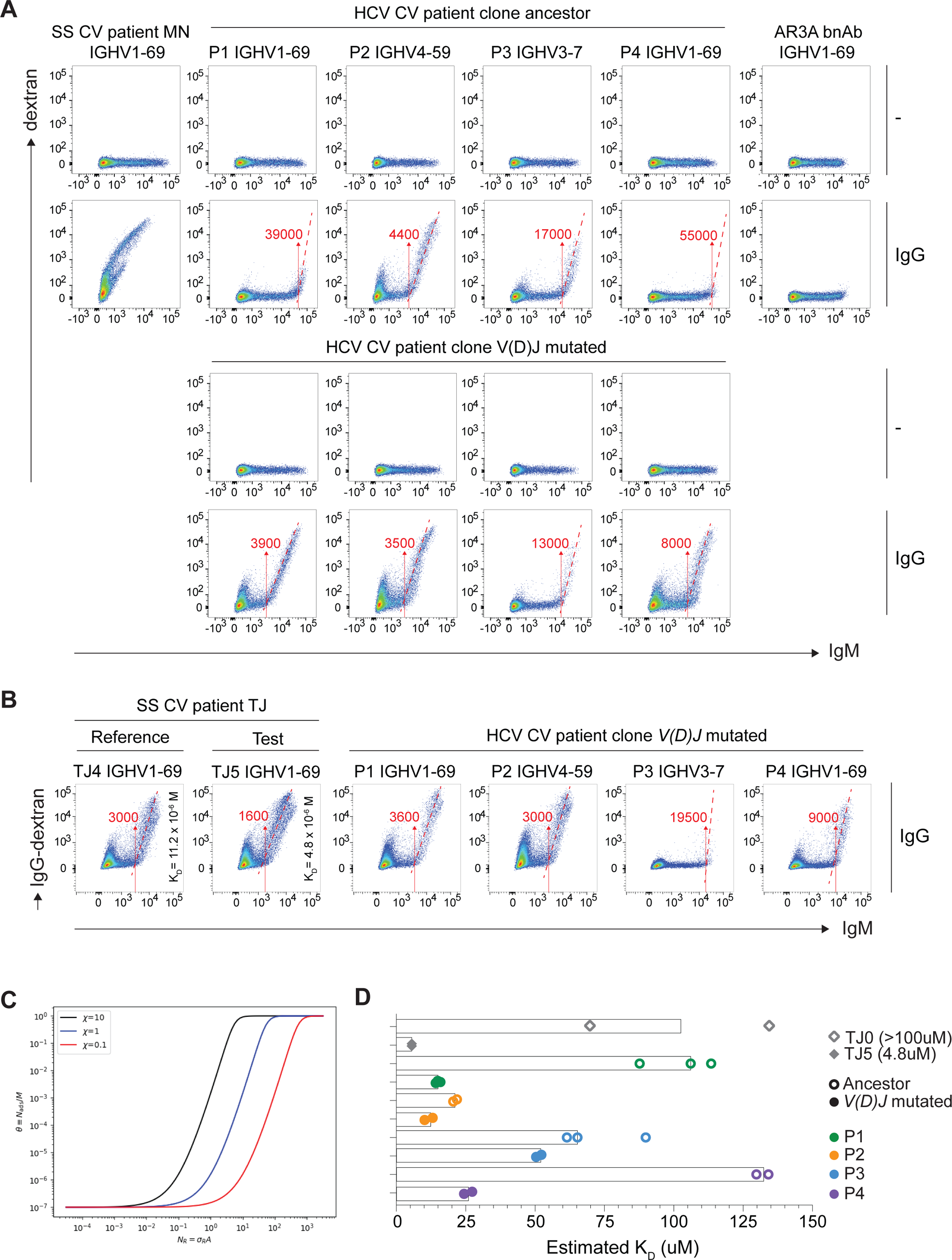
Estimating antibody affinities to IgG. A. Binding of empty dextran or IgG-dextran to HEK293 cells expressing membrane IgM corresponding to the B cell clone ancestor or the predominant *V(D)J* mutated clone from patients P1-P4. Control cells (far left and right columns) express membrane IgM corresponding to a mutated IGHV1-69/IGKV3-20 RhF cryoglobulin from a Sjogrens’s syndrome patient with vasculitis (patient MN) or a mutated IGHV1-69/IGKV3-20 broadly neutralising HCV E2 antibody (AR3A). HEK293 cells were transiently transfected and stained in parallel. The membrane IgM fluorescence intensity threshold for IgG-dextran binding is shown by the number and arrow in red. B. Comparison of IgG-dextran binding by cells expressing V(D)J mutated IgM from P1-P4 with cells expressing membrane IgM corresponding to two clonally related IGHV1-69/IGKV3-20 RhF autoantibodies with known affinities (TJ4 and TJ5). C. Mathematical modelling of absorption curves (e.g. fluorescence intensity, represented by 8) and surface receptor density (α*_r_*) for different bond strengths (ξ) (or affinities) for a fixed number of ligands (*N_L_*). The model shows the effect of increasing (or decreasing) bond strength (e.g. ξ =0.1; red, ξ = 1.0; blue and ξ = 10; black) causing the binding curve to rigidly shift to the left (or right). This relationship can be described by the equation described in the methods, Eq.5. D. Estimated affinity constants (K_D_) for IgG of each patient’s ancestor and mutated antibodies using Eq5 and IgM fluorescence intensities from 2-3 independent experiments, including the experiment shown in (B). RhF antibody TJ4 with known K_D_, is used as the reference antibody. RhF antibodies TJ5 and TJ0 with known K_D_, are used as validation antibodies. Bars (**|**) represent the mean K_D_. Example calculations shown in Figure S7.

The results above revealed the unmutated ancestor IgM cryoglobulins bound IgG when it was directly multimerised on dextran, raising the question of whether or not cells expressing these IgM autoantibodies would recognise IgG bound to a multivalent array of virus antigen. Cells expressing the IgM ancestor antibodies displayed no detectable binding of fluorescently labelled monomeric IgG even at a concentration 30-fold greater than the concentration in the IgG-dextran mixtures (13uM versus 0.4uM respectively; Figure S6C). However, when 6.7uM soluble monomeric IgG1 corresponding to the anti-E2 broadly neutralising antibody AR3C was mixed with E2 virus protein directly multimerised on dextran, cells expressing the ancestral P2 and P3 IgM displayed similar binding to the IgG1-virus antigen complex as they did to directly multimerised IgG-dextran (Figure S6C, far right column). Thus, while the P2 and P3 ancestral autoantibodies had no detectable binding to the virus E2 antigen, they bound self IgG when it decorated multimers of viral antigen.

### Estimating the affinity of IgG self-reactivity at clonal initiation

Transfected cells expressing the unmutated IgM ancestor of each patient’s pathogenic clone bound IgG-dextran, albeit in each case binding only occurred at a certain threshold level of membrane IgM that differed between the patients. Above the threshold, binding of IgG dextran increased steeply with small increases in membrane IgM (Figure 5E, bottom row, Figure 6A, second row, Figure S7A). Similarly, cells expressing the hypermutated cryoglobulin antibodies from each of the HCV-CV patients also bound IgG-dextran, and again, binding only occurred on cells above a threshold level of membrane IgM (Figure 6A,B). By contrast, cells expressing an unusually potent cryoglobulin from a Sjogren’s syndrome vasculitis patient, MN^45^, bound IgG-dextran in amounts linearly proportional to the amount of membrane IgM on each cell (Figure 6A, far left column).

The unmutated ancestor with the lowest IgM threshold for IgG-dextran binding, from patient P2, was the only ancestral antibody with detectable cryoglobulin activity (Figure 4B). This suggested that differences in the threshold amount of membrane IgM required for binding IgG-dextran might reflect differences in affinity. To test this, we transfected parallel sets of cells to express membrane IgM with affinities that had previously been determined as Fab’s by biolayer interferometry^45^. The mutated IGHV1-69/IGKV3-20 cryoglobulin, TJ5, and its immediate clonal precursor lacking one replacement mutation, TJ4, bind IgG with monovalent K_D_ of 4.8 x 10^−6^ M and 11.2 x 10^−6^ M, respectively. Consistent with a 2.3-fold higher K_D_, the threshold IgM per cell needed to bind IgG-dextran was 1.9-fold higher for TJ4 than for TJ5 (Figure 6B).

The shapes of the IgG-dextran binding curves, with a threshold level of membrane IgM required to observe binding, followed by a steep increase in binding above that threshold, correspond to the superselective binding reaction predicted by mathematical modelling of low affinity interactions between multivalent ligand in solution and surfaces bearing cognate receptors^79–83^ (Methods). The model predicts that, holding other variables constant, an increase in antibody K_D_ shifts the threshold amount of membrane IgM to a proportionally higher value (Figure 6C and Methods). As such, if the K_D_ value of a reference IgM is known, an unknown affinity constant (K_D_) can be extrapolated from the membrane IgM binding threshold and subsequent shift in the binding curve (Figure 6B, Figure S7B). To test the equation linking IgM threshold to K_D_ (Figure 6B, Figure S7C and Methods), we used the equation and threshold observed for TJ4 to calculate TJ5 IgG affinity to be K_D_ = 5.4 x 10^−6^ M (Figure 6B,D), and the unmutated ancestor of the TJ4 and TJ5 RhF antibodies, TJ0, to be K_D_ = 102 x 10^−6^ M (Figure 6D, Figure S7A), which were within range of their biolayer interferometry measurements of K_D_ = 4.8 x 10^−6^ M and K_D_ > 100 x 10^−6^ M respectively, indicating our method could extrapolate affinities within the micromolar range.

Hence, we used the equation to extrapolate the IgG affinities of the mutated pathogenic autoantibodies and their unmutated ancestors, using TJ4 as our reference (TJ4, K_D_ = 11.2 x 10^−6^ M). The affinities of the four V(D)J mutated pathogenic autoantibodies were estimated to be (from highest to lowest affinity): P2 K_D_ 11 x 10^−6^ M; P1 K_D_ 16 x 10^−6^ M; P4 K_D_ 26 x 10^−6^ M; and P3 K_D_ 51 x 10^−6^ M (Figure 6D). The estimated IgG affinities of the HCV-CV B cell clone ancestors were estimated to be (from highest to lowest): P2 K_D_ 21 x 10^−6^ M; P3 K_D_ 72 x 10^−6^ M; P1 K_D_ 100 x 10^−6^ M; P4 K_D_ 130 x 10^−6^ M (Figure 6D). It is problematic to measure binding affinities >10uM by most practical methods such as biolayer interferometry because it requires millimolar concentrations of Fab to be introduced to the sensor. The flow cytometric method developed here provides a practical way to quantify very low antibody binding affinities in the micromolar range corresponding to the first initiating interactions of an autoantibody response.

## Discussion

These findings address the question of why autoimmune CV arises in HCV-infected individuals, and why individuals with HCV-CV have a 35-fold increased risk of B cell lymphoma. In the four patients analysed in-depth here, three different sources of somatic mutation in B cells converged upon damaging combinations that allowed a pathogenic autoantibody-producing clone to accumulate in large numbers and cause clinical vasculitis. One source of somatic events was *V(D)J* recombination, a normal process in pre-B cells that fixed in the ancestral B cell and its progeny a particular IGHV/IGKV pair and a short CDR3_H_ length conferring low affinity binding to self-IgG, but not to HCV E2. The second somatic contributor to the disease was genome-wide accumulation of indel and SV/trisomy mutations, one or two of which conferred a growth or survival advantage on the self-reactive B cells, enabling their accumulation. The third pathogenic category was *V(D)J* hypermutation, which created clonal trajectories to produce insoluble IgM/IgG cryoglobulin complexes.

Out of hundreds of different IGHV/IGKV pairs frequently produced by *V(D)J* recombination, the pathogenic cryoglobulins in the four patients were specified by three IGHV/IGKV pairs that occur in >95% of HCV-induced cryoglobulins and monoclonal RhFs^24–26^. Antigenic mimicry between HCV and IgG could hypothetically explain this bias, since the IGHV1-69/IGKV3-20 pair found in 60% of cryoglobulins is also frequently used by HCV-neutralising IgG antibodies against the virus E2 protein. However, this hypothesis is not supported by the evidence here, since the IgM ancestor of the cryoglobulin in each patient bound self IgG but not E2, and the IGHV1-69/IGKV3-20 pairs that bind HCV have a longer CDR3_H_ than the pairs that bind IgG. Crystallographic studies of HCV-neutralising IGHV1-69 antibodies bound to E2 show that 40-60% of antibody buried surface area derives from CDR3_H_^19,22,23^. By contrast, crystallography of an IgG-bound IGHV1-69/IGKV3-20 antibody from a rheumatoid arthritis patient with macroglobulinemia showed CDR3_H_ contributes only 10% of buried surface area, and proposed that longer CDR3_H_ loops may prevent binding to IgG because they could no longer fit within the IgG C_H_2-C_H_3 cleft and would fill the binding pocket for a protruding IgG C_H_3 Leu432-His*cnf* loop^84^. The higher frequency of IGHV1-69/IGKV3-20 RhFs may simply reflect the higher frequency of this pair in the human B cell repertoire (5%) compared to the frequency of IGHV3-7/IGKV3-15 and IGHV4-59/IGKV3-15 pairs (1% and 2%, respectively)^85^.

Among thousands of somatic SNV, indel and SV mutations acquired genome-wide in each patient’s cryoglobulin-producing clone, only one or two per clone corresponded to putative driver mutations found recurrently in human B cell lymphomas and leukemias. Two complementary bodies of evidence support the conclusion that the specific nature of these mutations provided a growth or survival advantage explaining the large size of these clones. Firstly, the loss-of-function mutations in *KLF2, TRAF3* and *BIRC3*, and gain-of-function mutations in *NOTCH1* and chromosome 12, occur frequently in neoplastic B cell clones and in their asymptomatic precursor, MBL, but were not observed in normal memory B cells^51^. Secondly the mutated genes regulate key biochemical events in B cell growth and survival. The 11q22-23 deletion results in dual loss of *BIRC3* and *BIRC2*, while the *TRAF3* indel mutation creates a loss-of-function. TRAF3, BIRC3 and BIRC2 are critical negative regulators of the NF-κB signalling pathway that mediates the response to B cell growth and survival factors CD40L, BAFF or TLR ligands^65,69,86^. The *KLF2* mutation diminished its activity as a negative regulator of NF-κB. *KLF2* dampens expression of NF-κB-dependent genes encoding CD21 and CD23, and enforced *KLF2* expression in LPS-activated B cells inhibits expression of the NF-κB-induced gene *IRF4*^87–89^. The gain-of-function *NOTCH1* mutation prevents degradation of the intracellular NOTCH fragment, a fragment produced in B cells contacting lymph node stroma and known to promote expression of B cell proliferation genes cyclin D and *MYC*^90,91^. Trisomy 12 increased expression of chromosome 12 genes in the clonal B cells. Among ch12 genes are drivers of cell cycle (*CCND2*, *CDKN1B*, *CDK2*, *CDK4*), B cell lymphomagenesis (*BCL7A*, *P2RX7*, *BTG1*, *KRAS*, *KMT2D*, *STAT6*), and B cell signalling (*PIK3C2G*, *IRAK4*). The acquisition of lymphoma driver mutations in the autoimmune clones provides an explanation for the 35-fold higher incidence of B cell lymphoma in HCV patients with CV compared to the 2.4-fold increased incidence in HCV patients overall^36,37^. Current international guidelines do not provide specific advice on monitoring HCV infected or HCV-CV patients for the development of lymphoproliferative disease after DAA treatment (https://www.who.int/publications/i/item/9789240052734). Considering the presence of driver mutations and the persistence of the cryoglobulin-producing B cell clones after DAA therapy and viral clearance, it highlights the need for longitudinal studies to test if patients remain at elevated risk of B cell lymphoma.

The genome of memory B cells in healthy donors progressively accumulates SNV and indel mutations throughout life, with a mean of 2194 SNVs and 94 indels in people over 50 originating in ageing hematopoietic stem cells and during naïve B cell proliferation and differentiation into memory B cells^51^. Since the HCV-CV clones had comparable genome-wide mutation burdens to memory B cells from healthy older donors, their clonal expansion does not appear to reflect increased genome-wide mutagenesis but instead one or two of the indels or SVs in the pathogenic memory B cells happened to occur in genomic locations that create a growth or survival advantage. Conversely, the HCV-CV clones had fewer SVs than CLLs, potentially explaining why the autoantibody-producing clones do not accumulate to greater numbers.

The findings here about starting *V(D)J* specificity and genome-wide mutation burden inform a specific hypothesis to explain why HCV so frequently creates a perfect mutation storm in a single B cell. Plasma IgG is effectively monovalent for binding to “public idiotype” monoclonal RhFs triggered by chronic HCV^92–94^. While the plasma IgG concentration (10^−4^ M) is above the K_D_ of these antibodies, it may not crosslink membrane IgM receptors to trigger B cell tolerance checkpoints. Transgenic mouse experiments tracing B cells displaying membrane IgM with low-affinity for self IgG (AM14, K_D_ = 2 x 10^−6^ M) have shown circulating plasma IgG does not trigger B cell tolerance checkpoints in these cells^95^. However when a small amount of anti-DNA or anti-RNA IgG is multimerised on nucleic acid-containing apoptotic cell nanoparticles, it acquires the capacity to bind stably to the AM14 IgM-expressing B cells measured by flow cytometry, and to stimulate their proliferation, *V(D)J* hypermutation and RhF secretion^96–98^. Similarly, we showed here that multimerization of anti-HCV IgG on HCV E2-dextran conferred flow cytometrically detectable binding to cells expressing membrane IgM corresponding to public idiotype RhFs.

In chronic HCV infection, large amounts of viral RNA circulates in plasma primarily within IgG-bound 35-65 nm particles^99–101^. Over time, these circulating IgG-virus nanoparticles may activate large numbers of naïve B cells emerging from *V(D)J* recombination with public idiotype RhFs, promoting each into proliferation and further somatic mutagenesis. The result is a perfect storm where the normal processes of genome-wide mutagenesis, *V(D)J* recombination and *V(D)J* hypermutation converge by chance on rare combinations that produce an autoantibody in sufficient quantity and quality to trigger clinical vasculitis in 5-15% of infected individuals.

In summary, this work reveals that one of the strongest associations between virus infection and autoimmune disease results from a pathogenic confluence of multiple somatic mutation events within a self-reactive B cell clone. Given that each of the sources of somatic mutation are common to all memory B cells, it is conceivable that similar processes contribute to the pathogenesis of other autoimmune diseases where the environmental trigger is less understood.

### Study limitations

First, the study is limited to four HCV-CV patients. Future studies of larger numbers of B cell clones will be needed to determine the prevalence of the findings here for HCV-CV, and to determine to what extent the somatic variant burden in the pathogenic clones from HCV-CV patients differ from those in normal memory B cells, malignant B cells, or in self-reactive memory B cells clones in other autoimmune diseases. It is also unclear if the same somatic mutation burden would be identified in patients with HCV infection and a detectable cryoglobulin RhF, but without clinical vasculitis. Second, we have not directly demonstrated the individual driver mutations present in each patient B cell clone provides a growth or survival advantage *in vivo*. Third, whilst we did not detect binding of ancestor antibodies to the HCV E2 protein, this does not exclude the possibility of a very low affinity interaction with E2 below the sensitivity of the multimerised dextran flow cytometry assay, nor with other HCV proteins, so antigenic mimicry with HCV cannot be conclusively excluded.

## Supporting information

Table S1 Patient clinical data

Table S2 P1 clone vs polyclonal memory

Table S3 SNV and indels

Table S5 Ploidy

Table S6 Lymphoma genes

Table S7 Primers

Table S8 Antibodies

## Acknowledgments

We thank the entire Goodnow laboratory, Daniel Christ and David Langley for insightful discussions. We thank Daniel Lingwood for membrane IgM assay advice and discussion, and Heather Machado for advice on somatic mutations in memory B cells. We thank Shane Grey for gifting the pcDNA3.1 empty plasmid and Daniele Cultrone for technical assistance with the NF-κB luciferase assay. We thank the Garvan-Weizmann Centre for Cellular Genomics, the Kinghorn Cancer Centre for Clinical Genomics, the Garvan Sequencing Platform and the Ramaciotti Centre UNSW Sydney for technical services. This work was supported by the Bill and Patricia Ritchie Foundation, the Croall Foundation, National Health and Medical Research Council Australia (NHMRC) grants APP2010134 & APP1113904, UNSW Cellular Genomics Futures Institute and a UNSW Scientia Postgraduate Scholarship.

## Author contributions

Patient selection and recruitment M.D, A.C, A.D.K, G.J.D, G.M and D.S, flow cytometry and analysis C.Y, antibody mRNA deep sequencing and analysis C.Y, K.J.L.J, M.L.F, J.H.R, single-cell RNA sequencing and analysis C.Y, M.S, K.J.L.J, T.J.P, M.G, G.A, F.L, whole genome sequencing and analysis C.Y, M.S, M.A.F, K.J.L.J, T.J.P, S.R and C.C.G, single-cell DNA sequencing and variant validation including luciferase assay C.Y and M.S, antibody ancestor reversions C.Y, K.J.L.J, antibody production and binding/cryoaggregation assay C.Y, M.L.F, J.H.R, antibody peptide sequence analysis J.J.W and T.P.G, *in vitro* membrane antibody assay C.Y, M.L.F, D.A, R.A.B, CDR3 length analysis K.J.L.J and C.C.G, mathematical modelling S.A-U, D.F, visualisation C.Y, M.S., K.J.L.J, T.J.P, S.A-U, supervision R.B, D.S, C.C.G, study design and writing, C.Y, D.S, C.C.G. All authors contributed to manuscript edits.

## Methods

### Patient samples

Peripheral blood was taken from patients with chronic Hepatitis C virus (HCV) infection preparing to undergo direct acting antiviral (DAA) treatment recruited under the Surveillance for Antiviral Resistant Variants in Chronic Hepatitis C (SEARCH-C) study, St Vincent’s Hospital (SVH), Sydney, Australia. The SEARCH-C was approved by the St Vincent’s Hospital Human Research Ethics Committee (HREC/11/SVH/197). Peripheral blood was collected and processed by Ficoll-Paque centrifugation for peripheral blood mononuclear cells (PBMC) collection. Blood samples were collected before the start of DAA therapy (baseline) and again between 25-50 weeks later. At the baseline timepoint, the patients commenced 12 weeks of DAA therapy. All patients achieved HCV clearance which was confirmed by absence of HCV RNA in serum. Four HCV patients from the SEARCH-C study were retrospectively identified as having been diagnosed with HCV-associated cryoglobulinemic vasculitis before recruitment into the SEARCH-C study based on their clinical features, laboratory and biopsy results. Patient P3 suffered from glomerulonephritis and was treated with rituximab 7 weeks before the baseline sample (HCV^+^) and commencement of DAA therapy, and patient P4 experienced a more severe neuropathy and was treated with methotrexate, mycophenolate and prednisolone both before and after DAA (Table S1). All patients were consented and their samples were transferred to and analysed under the SOMAD study/HOPE Research Program at the Garvan Institute of Medical Research, Sydney, Australia approved by the Western Sydney Local Health District Human Research Ethics Committee (HREC17/5449).

### Flow cytometry

PBMCs were thawed and washed in 2% FCS/PBS and incubated with Fc block for 20 min on ice. PBMCs were stained with eF780 Fixable Viability Dye (eBioscience) and a cocktail of antibodies (Table S8) for 30 mins on ice. Anti-idiotypic monoclonal antibody G6, specific for F-allele IGHV1-69 heavy chains, was fluorescently labelled using the Alexa Fluor 488 Antibody Labelling kit (Life Technologies)^102^. After staining, cells were washed and fixed in 2% formaldehyde (5% formalin, Sigma) for 15 min on ice.

### Fluorescence activated cell sorting (FACS)

Cells were either bulk sorted for WGS or single cell RNA-sequencing (10X Genomics) or single cell sorted for downstream single cell RNA-sequencing (Smart-seq2), either done alone or in parallel with single cell DNA sequencing (G&T). For both bulk and single cell sorting, PBMCs were stained using the same buffers and steps as described for flow cytometry (above), with the exception of the anti-idiotypic monoclonal antibody G6, which was only used to stain the PBMCs of patients P1 and P4. For bulk sorting for whole genome sequencing, two populations from the baseline timepoint were sorted: IgM^+^ B cell clone and polyclonal B cells, according to the gating strategy shown (Figure 2B). For single cell sorting for immunoglobulin and DNA analysis, total B cells (CD19^+^CD20^+^) or memory B cells (CD27^+^) from the baseline (HCV^+^) timepoint were sorted into 96 well Lo bind plates containing buffers as described below. For single cell RNA-sequencing using the 10X Genomics platform, total B cells (CD19^+^CD20^+^) from patient P1 PBMC were sorted at the clearance (HCV^−^) timepoint.

### Bulk immunoglobulin sequencing

Total RNA from roughly 1-3 million thawed patient PBMCs was extracted using a RNeasy Mini Extraction kit (Qiagen) and reverse transcribed to cDNA using oligo-DT primers and the iScript cDNA Synthesis Kit (Bio-Rad). Immunoglobulin heavy and light chain cDNA was amplified using the Q5 High-Fidelity 2X Master mix (NEB) and pooled forward primers that bind the leader sequences of the variable regions and the reverse primers that bind the first exon of the mu and kappa constant regions, as previously described^45^ (Table S7). The primers incorporate 5’ sequences compatible with sequencing on the Illumina platform. Two separate PCR reactions were performed for each antibody isotype i.e. mu and kappa. PCR products were indexed (Illumina), pooled and sequenced on an MiSeq machine (Illumina) capturing 300bp paired end reads to a depth of 1 million reads per sample. B cell clones were identified using the MiXCR software tool (v3.0.12) (https://github.com/milaboratory/mixcr.git)^103^, with the top clones based on total number of reads sharing >90% CDR3 amino acid identity. The top five clones were displayed as proportion of all sequenced reads.

### Single cell RNA sequencing

Libraries for single cell RNA sequencing on the patient baseline samples (HCV^+^) were prepared using Smart-seq2 protocol^104^. If genomic DNA was also being extracted simultaneously, cells were processed as described for the Genome & Transcriptome (G&T) protocol^105^. The Smart-seq2 protocol was modified to halve the volume of lysis buffer added to each well of the plate (from 4 µL to 2µL), ensuring the lysis buffer/oligo-DT/dNTP was maintained at a ratio of 2:1:1. cDNA was generated as described^104^, but with reactions performed at half the volumes. cDNA was amplified as described^104^ but with the ISPCR primer final concentration reduced to 50nM. For each cell, 1ng of DNA was added to a new 96 well plate for library generation. Amplified DNA was tagmented and barcoded using the Nextera XT DNA Library Preparation Kit (Illumina). The same quantity of each barcoded DNA libraries (range 4-10ng) were pooled together for sequencing. DNA libraries were sequenced on an NextSeq550 machine (Illumina) at a median depth of 1 million reads per cell.

Single cell V(D)J and RNA-sequencing at the clearance (HCV^−^) timepoint for blood of patient P1 was performed using Repertoire and Gene Expression by Sequencing (RAGE-Seq) using droplet based capture of 10,000 cells using 10X Genomics Chromium 3’ system as described^48^. In brief, following reverse transcription and before fragmentation, two extra PCR cycles were performed and full length cDNA equally spilt for short-read gene expression library generation (Illumina) and targeted V(D)J capture followed by long-read library generation (Oxford Nanopore Technologies, ONT). Short-read gene expression libraries were sequenced on a NovaSeq 500 (Illumina) at 50,000 mean reads per cell. Long-read V(D)J libraries were loaded onto R9.5.1 (FLO-MIN107) flowcells (ONT) with base calling performed using the Albacore software pipeline (v2.2.7) (ONT).

### Single cell immunoglobulin analysis and clonal tree generation

The immunoglobulin B cell receptor (BCR) heavy and light chain nucleotide sequences for each cell of patient P1 clearance (HCV^−^) sample were generated by targeted capture and long-read sequencing using the RAGE-Seq protocol, were generated as described^48^. In brief, long-read sequencing data was demultiplexed by cell barcode, subject to *de novo* assembly, aligned and “polished” to generate fasta files containing consensus transcript contigs. The immunoglobulin sequences were determined by aligning the contigs to IgBLAST and BLASTN to determine *V(D)J* and constant region gene usage respectively. Immunoglobulin sequences that were non-productive, out-of-frame or contained stop codons were removed. Each cell was annotated as polyclonal or clonal, where clones were defined as having >90% amino acid identity with the heavy chain and light chain CDR3 of P1 B cell clone (Figure 1E).

The immunoglobulin B cell receptor (BCR) heavy and light chain nucleotide sequences for each cell generated by the Smart-seq2 protocol from the patient baseline (HCV^+^) samples was reconstructed from the single cell RNA-seq data using the VDJPuzzle software package (https://bitbucket.org/kirbyvisp/vdjpuzzle2)^106^. The individual heavy and light chain nucleotide sequences obtained by VDJPuzzle were uploaded to IMGT HighVQuest (v1.9.1). Using the somatic hypermutation annotations, a clonal tree was constructed. The diameter of each circle is proportion to the number of individual B cells sharing an identical immunoglobulin sequence, with the exception of the unmutated ancestor circle (grey). The joining lines between the circles present number of somatic hypermutations between the immunoglobulin sequences, where the length of the lines is proportional to the number of somatic hypermutations (linear scale, 1 unit of distance = 1 mutation). Somatic *V(D)J* mutations over the whole *V(D)J* gene loci were counted i.e. including CDR3 and FR4.

### Single cell RNA seq gene expression analysis

Raw sequencing files binary base call (bcl) files were demultiplexed and converted to Fastq using bcl2fastq (v2.19.0.316) (Illumina). Alignment and barcode counting were performed using cell ranger (v6.0.2) (10X Genomics). Gene expression reads were aligned to human genome reference GRCh38 (hg38). Gene expression count matrices were exported by cell barcode linked to a polyclonal or clonal annotation (as determined above) using R. Single cell gene expression counts were normalized using SAVER with default values^107^. Single cells were excluded if the library size or number of expressed genes fell below 2 median absolute deviations, or if mitochondrial reads accounted for more than 20% of the reads. Polyclonal naïve and B cell memory clusters were determined using landmark genes *IGHM*, *IGHG1*, *IGHD*, *TCL1A.* Differentially expressed genes were determined using limma on log normalised values^108^. Bonferroni correction was applied to each set of p-values, to determine the family-wise error rate (FWER). Statistically significant differentially expressed genes were defined as FWER <0.05.

### Whole genome sequencing

For the two patients with an IGHV1-69 B cell clone (P1 and P4), the G6 antibody was used to sort the expanded B cell clone (G6^+^ IgK^+^ B cells) and the control polyclonal B cell population (G6^−^ B cells) (Figure 2B). For the other two patient B cell clones encoded by heavy chains for which no anti-idiotypic antibody exists (P2; IGHV4-59 and P3; IGHV3-7), the B cell clone was identified and sorted from the IgM^+^ kappa light chain restricted memory B cell population (CD27^+^ IgM^+^ IgK^+^ B cells) and the control polyclonal B cell population consisted of all remaining B cells that did not co-express IgM and IgK (Figure 2B). These gating strategies were predicted to result in purity >90% (>95% for the IGHV1-69 B cell clones).

Approximately 30,000 cells (minimum 8,000) of either the B cell clone or polyclonal B cells of the patients were directly FACS sorted into 100µL ALT lysis buffer (Qiagen MicroKit). gDNA was extracted as per manufacturer’s instructions (Qiagen MicroKit). Approximately 2/3 of gDNA was taken to generate DNA libraries (minimum 10ng), with remaining gDNA to be stored for variant confirmation. DNA libraries for whole genome sequencing were generated using the Kapa Hyper Plus Kit (Roche) for low input DNA. DNA libraries were generated according to manufacturer instructions, with the following modifications: fragmentation for 17 minutes, adapter ligation for 1 hour and PCR amplification with 3-5 cycles depending on starting DNA amount (5 cycles for ∼10ng DNA). DNA libraries were sequenced on a HiSeqX instrument (Illumina) with 150bp paired end reads at 30X coverage for the polyclonal population and at 60X coverage for the B cell clone. WGS sequencing quality was verified by calculating the coverage (unique reads) per nucleotide base over ∼180 genes at exonic locations totalling ∼1,430kb, using the BAM files generated from somatic variant analysis and BEDtools.

### SNV mutational signatures

To detect small somatic variants (SNV), the Dynamic Read Analysis for Genomics (DRAGEN) Somatic Pipeline (v3.3.7) (Illumina) for tumour/normal pair was used, with the expanded B cell clone assigned as “tumour” and polyclonal B cells assigned as “normal.” Reads were aligned to GRCh37 (hg19) reference genome. The COSMIC (v3) single base substitution (SBS) mutational signatures of somatic SNV calls of the HCV CV clones were fit to mutational signatures relevant to lymphocytes (SBSBlood, SBS1, SBS7a, SBS8, SBS9, SBS18) as described^51^ using the R package sigfit (v2.0)^109^. The SBS signatures were plotted as a relative proportion for each clone.

### Small (<50bp) somatic variant (SNV and indel) analysis

Four PCAWG tumour/normal sample pairs from the CLL-mutated (“Lymph-CLL”) and BNHL (“Lymph-BNHL) patient cohort were identified^49^ and extracted from ICGC Controlled Data. Variants were called using a workflow previously described^110,111^. In brief, the PCAWG BAM files were downloaded and HCV B cell clone tumour/normal BAM files generated using the Dynamic Read Analysis for Genomics (DRAGEN) Somatic Pipeline (v3.3.7) (Illumina) (see above) were used to extract reads using SAMtools^112^ bam2fq. Read were aligned to GRCh38.p14 (hg38) using BWA^113^ and prepared for variant calling using GATK best practices^114^. For detecting somatic SNV and indel calls, default filtering was applied for both GATK-MuTect2^114^ and Strekla2^115^ and for each sample a consensus of SNVs and indels were generated using an approach previously described^116^ (Table S3). Somatic variant output file was annotated using the Ensembl Variant Effect Predictor (VEP) algorithm^117^. Somatic variants were filtered selecting for coding variants resulting a frameshift, in-frame deletion or insertion, missense, stop gain or loss or synonymous variant that occurred within a gene of a curated list of 221 genes frequently mutated in B cell lymphomas (Table S6). Somatic variants were verified using the Integrative Genomics Viewer (IGV) (v2.5.3) and used to calculate the proportion of reads carrying the variant (i.e. variant allele frequency, VAF).

### Large (>50bp) structural somatic variant analysis

To detect somatic SVs in the patient HCV CV clones, the CLL-mutated (“Lymph-CLL”) and BNHL (“Lymph-BNHL) samples (see above)^49^, Manta^118^ and Delly^119^ were run in somatic tumour/normal mode with default filtering. Manta and Delly identify and score possible SVs based on two types of evidence; discordant pair end reads and split read alignments. For each sample, a consensus of SVs were generated using an approach previously described^116^ (Table S4). The consensus SV list was manually annotated for variants occurring at the heavy, kappa or lambda immunoglobulin loci using the hg38 UCSC Genome Browser.

### Somatic ploidy analysis

To detect the presence of any ploidy event such as a chromosome gain, the average whole genome sequencing coverage was calculated for the whole genome for chromosomes 1 to 22 using the Samtools depth command^62^. An average across every base for the entire genome for each of the B cell clone and the polyclonal B cells was first calculated, and this ratio utilised as a baseline. The same was done for each chromosome and this ratio compared to the whole genome baseline ratio (Table S5). If a whole chromosome gain was present, we would expect ∼1.5X more reads in the chromosome ratio than the baseline genomic ratio. Any chromosome loss (monoploidy) was defined as a ratio ≤0.6, any single whole chromosome gain (triploidy) was defined as a ratio ≥1.4 and whole duplication of both chromosomes (tetraploidy) was defined as a ratio ≥1.9.

### Somatic variant validation

PCR with or without sanger sequencing was performed on either remaining bulk sorted gDNA or from single cell amplified gDNA performed in parallel with mRNA extraction for single cell immunoglobulin sequencing using the Genome & Transcriptome (G&T) protocol^105^. Following separation of single cell mRNA and gDNA, gDNA was amplified via multiple displacement amplification (MDA) using the REPLI-g Single Cell Kit (Qiagen) according to manufacturer’s instructions. MDA rapidly amplifies DNA at a constant temperature, by using a combination of random hexamer primers and a high-fidelity bacteriophage DNA polymerase (error rate 1 in 10^6^-10^7^ bases). PCR primers and PCR conditions used on bulk or single cell derived gDNA are as indicated (Table S7). All PCR primers were designed in house with the exception of the KLF2 primers^54^. All PCR reactions used a standard Taq polymerase (Invitrogen), with the exception of KLF2 which was performed using the Q5 High-Fidelity kit (NEB). For the 11q deletion variant (P3), primers for a control gene (*SEC23IP*) giving rise to a PCR product half the size was also included in the same PCR reaction. Purified PCR products (with the exception of the *SEC23IP* control gene) were submitted for Sanger sequencing.

### NF-κB luciferase assay

Adherent HEK293T cells were co-transfected with 0.3μg NF-kB luciferase reporter (pcDNA3.1-luc) (Promega), 0.2μg of β-galactosidase plasmid (CMV.β-galactosidase, gifted by Shane Grey) and either 0.5μg pcDNA3.1 vectors encoding wildtype or KLF2 variants or empty pcDNA3.1 vector (total 1μg DNA). Transfection was performed using Lipofectamine 3000 (ThermoFisher) in serum-free conditions. Two hours before harvest, cells were stimulated with human recombinant TNFα (R&D Systems) at 300IU/mL. Luciferase activity was measured in cell lysates collected 8 hours after transfection (Promega). To correct for transfection efficiency, luciferase results were normalised to β-galactosidase activity measured (Galacto-StarTM).

### Soluble IgM antibody expression

Secretory IgM antibodies were produced in house by transient transfection of non-adherent Expi293F (Gibco) cells using the Expi293 Expression System Kit (Gibco). The heavy immunoglobulin variable region sequences were inserted into the human heavy chain mu constant region secretory sequence (Genbank accession number BC073758.1) and synthesised into mammalian vector pcDNA3.1 (Genscript). Similarly, the light chain immunoglobulin variable region sequences were inserted into the human kappa constant region sequence (Genbank accession number OM584289.1) and synthesised into mammalian vector pcDNA3.1 (Genscript). Human J chain was also synthesised into mammalian vector pcDNA3.1. Expi293F cells were transfected according to manufacturer’s instructions with a 10µg of heavy chain plasmid, 10µg of light chain plasmid and 10µg of J chain plasmid (ratio 1:1:1) for total of 30µg DNA per flask at a density of ∼3 x 10^6^ cells/mL. On Day 7 post-transfection, supernatant for each flask was collected, and spun for 10 min at 3000g and passed through a 0.22µM filter. Soluble IgM was purified by concentrating the supernatant using centrifugal filter units (Amicon, Merek) with a 30K membrane cut-off spun at 4000g for 40-60 mins at room temperature until <2mL concentrate remained.

### Soluble IgM quantification by ELISA

A 384-well plate (Corning) was coated with 0.5µg/mL purified anti-human IgM diluted in 1X PBS for 1 hour at room temperature and blocked with 3% BSA/1XPBS for 1 hour at 37°C. An IgM standard was prepared using human whole IgM derived from serum (Jackson ImmunoResearch) starting at 10µg/mL concentration followed by a 1:2 serial dilution. Concentrated supernatants containing soluble IgM were diluted 1:100 followed by a 1:2 serial dilution performed in duplicates and incubated for 1 hour at room temperature. Presence of IgM was detected using a goat anti-human IgM (Fc5u specific) secondary antibody conjugated to alkaline phosphatase (Jackson ImmunoResearch) (1:1000) incubated for 1 hour at room temperature before addition of para-nitrophenylphosphate substrate (Sigma Aldrich). ELISA plate was read an optical density (OD) at 405nm (ClarioStar instrument, BMG Labtech). Concentration of IgM (µg/mL) in each supernatant was interpolated using the standard curve.

### IgG (RhF) ELISA

384-well plates (Corning) were coated with either 5µg/mL whole IgG obtained from human serum (Jackson ImmunoResearch) in NPP coating buffer overnight. ELISA plates were blocked with 3% BSA/PBS for 1 hour at 37°C. Samples were added in duplicate or triplicate to the plate at a starting concentration of 100µg/ml, followed by 1:2 serial dilution and incubated for 1 hour at 37°C. Anti-human IgM-alkaline phosphatase (Sigma Aldrich) (1:1000) was used as secondary detection antibody incubated at 45 mins at 37°C before addition of para-nitrophenylphosphate substrate (Sigma Aldrich). Plates were read at an optical density (OD) at 405nm (ClarioStar instrument, BMG Labtech).

### Cryoglobulin assay

Cryoglobulin activity was analysed via a temperature-controlled spectrophotometer (Cary Series UV-Vis spectrometer, Agilent). Antibody cryoaggregation causes changes in light scattering^120^. IgM supernatants stored at 4°C were allowed to warm to 37°C on a heat-block for 20 min (cryoglobulins redissolve upon warming). An equal volume of IgM antibody at a concentration of 150-200µg/mL was mixed with equal volume of human IgG obtained from blood (Sigma-Aldrich) at a concentration of 1mg/mL (ratio ∼1:5) in 1X PBS. The mixture of IgM and IgG was incubated for another 20 min at 37°C on a heat-block before immediately transferring to the spectrometer set to 37°C. Approximately 200 optical density (OD) measurements were taken at 500nm as the spectrophotometer reduced in temperature by 0.1°C/min from 37°C to 4°C.

### RhF peptide sequencing and data analysis

IgM RhF antibody was purified from patient P3 plasma by precipitating with heat-aggregated IgG followed by reduced SDS-PAGE^121^ and non-reduced SDS-PAGE^45^, respectively. The IgM RhF gel band of the heavy and light chain was excised and digested with Pierce trypsin protease (ThermoFisher Scientific) and chymotrypsin (Promega) to generate peptides for LC-MS/MS using a Thermo Scientific Orbitrap Exploris 480 mass spectrometer coupled to an Ultimate 3000 UHPLC (Dionex). Peptide sequence analysis was performed by de novo sequencing and the IMGT database matching using Peaks studio XPro software (Bioinformatics Solution Inc., Waterloo, ON, Canada). Parameters for database searches, data refinement and immunoglobulin variable region subfamily assignments were described previously^121,122^. An average local confidence score threshold of 80% was applied to select high quality de novo peptides. A false discovery rate (FDR) threshold of 1.0% was applied at the peptide level to each data set. The immunoglobulin variable region subfamily is assigned from the presence of a unique peptide corresponding to the subfamily.

### Recombinant HCV E2 (including biotinylated)

The E2 protein sequence was derived from HCV Genotype 1a (isolate H77, GenBank accession number AF011751) and HCV Genotype 3a (isolate UNK3a.13.6, GenBank accession number AY894683). Recombinant HCV E2 (rE2) antigen was produced as described^123^. In brief, the E2 construct was incorporated the HCV polyprotein amino acid residues 384 to 661 encoding the E2 ectodomain (no stem or transmembrane domain) into a pcDNA3.1 vector with a N-terminus secretion signal, a C-terminal Avitag™ and a six-histidine tag (6×His) and transiently transfected into HEK293F cells. Media was harvested 96 hours later and rE2 was purified by passing through a 1 ml HiTrap chelating HP chromatography column. Biotinylation of rE2 was performed at the C-terminal Avitag™ using E.coli biotin ligase (BirA).

### Surface IgM expression and antigen staining

Methods for transient transfection for expression of membrane-bound IgM on ExpiHEK293F cells were adapted from the published protocol^77^. Desired heavy immunoglobulin variable region sequences were inserted into the human heavy chain mu constant region membrane sequence that includes the transmembrane region (Genbank accession no. BC009851.2) and synthesised into mammalian vector pcDNA3.1 (Genscript). Immunoglobulin light chain constructs were the same as used for secretory IgM expression. Expi293F cells were transfected according to manufacturer’s instructions with 10µg of heavy chain plasmid and10µg of light chain plasmid (ratio 1:1) for total of 20µg DNA per flask at a density of approximately 1.2 x 10^6^ cells/mL. On Day 2 or Day 3 post-transfection, Expi293 cells were harvested and approximately 0.5 x 10^6^ cells were transferred to a 96 well U bottom plate (Corning). Cells were stained with 5uL of biotinylated IgG (2mg/mL, Jackson Immunoresearch) or 5uL of commercial fluorescently labelled dextran molecule (Klickmer^®^, Immudex) loaded with biotinylated IgG, biotinylated rE2 or biotinylated rE2 bound with anti-E2 monoclonal antibody, for 10 min at room temperature. After staining with the antigen loaded Klickmer dextran, anti-human IgM antibody, anti-human IgK antibody and live/dead viability dye (Table S8) was added to the cells and incubated for a further 25 min on ice. Cells were washed in 1XPBS by removing supernatant from cell pellets by pipetting. Cells were run immediately on a LSRII Fortessa or Symphony (BD) instrument. To obtain the IgM fluorescence intensity at which antigen binding occurs, the geometric mean fluorescence on a narrow gate of ∼100 events was generated at the IgM antigen binding inflection point as identified by visual inspection using FlowJo^TM^ software (v.10).

Klickmer^®^ molecules carried 25 or 20 biotin acceptor sites attached along a dextran backbone containing phycoerythrin (PE). Biotinylated E2 (described above) or biotin-SP (long spacer) ChromPure human IgG (Jackson Immunoresearch) were mixed with PE-Klickmers according to manufacturer’s instructions (HCV rE2; 60kDa, whole IgG; 150kDa) at molar ratios to achieve full loading of acceptor sites. Antigen was incubated with Klickmer^®^ for 30 mins in the dark at room temperature. Where required, Klickmer^®^ molecules loaded with biotinylated rE2 were incubated with 5uL purified anti-E2 AR3C monoclonal IgG1 antibody (1mg/mL, Genscript) for 20 minutes in the dark. Antigen-Klickmer^®^ molecules were diluted with 1XPBS according to manufacturer’s instructions, ready for staining. Antigen-Klickmer^®^ mix was stored in the dark at 4°C for up to 12 weeks.

### IGHV1-69/IGKV3-20 anti-HCV E2 antibodies isolated from an HCV patient

Antibodies B04 and H04 were identified by single cell mRNA sequencing of single cell sorted HCV E2 (cocktail of Genotype 1a and 3a) tetramer-binding memory B cells from a patient chronically infected with HCV (Genotype 3a)^123^(Rowena A Bull, manuscript in preparation). Antibodies were confirmed to be HCV E2 specific by ELISA. Mutations in heavy and light chains were reverted to the unmutated ancestor using IMGT. Non-synonymous mutations in the heavy chain and light chain CDR3 were also reverted if the mutation resided within the V and J gene. In the case of H04 and B04, non-synonymous mutations were also reverted in the D genes (deemed to reside at positions clear of N addition sites).

### IGHV1-69 CDR3 amino acid lengths

Seventeen IGHV1-69/IGKV3-20 RhF antibody sequences confirmed to bind IgG were acquired from this project (P1,P4) and published data^33,34,45,84,124^. Ten IGHV1-69/IGKV3-20 anti-HCV E2 antibody sequences were derived from the same HCV patient as described above (Rowena A Bull, manuscript in preparation). IGHV1-69/IGKV3-20 antibody sequences with unknown specificity were obtained from single memory B cells from three healthy donors^85^. Unmutated (<2% SHM) IGHV1-69 IgM (“naïve”) and mutated (≥ 2% SHM) IGHV1-69 IgG (“memory”) antibody sequences were from bulk repertoire sequencing of healthy donor PBMCs filtered for least 50,000 clones (61 healthy adult donors)^125^. CDR3 amino acid lengths were defined as according to IMGT, with anchor residues removed.

### Mathematical modelling of multivalent antigen binding to IgM expressed on cells

Our mathematical model for surface IgM expression binding to IgG-dextran arises from adapting previous results on the interaction between multivalent ligand-functionalised nanoparticles and surfaces bearing cognate receptors^79–83^. Equilibrium binding probabilities in multivalent systems can be accurately described within the context of statistical mechanics. If a receptor coated surface (i.e. surface IgM expressed on HEK cells) is in contact with a solution containing ligand-functionalised constructs (i.e. IgG-dextran)^79^, the number of constructs bound is given by the Langmuir formula:

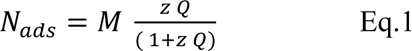

*M* is the maximum number of adsorption sites on the surface and *Q* = *Q*(*N_L_*, *σ_R_*, *χ*), the binding partition function, depends on the number of ligands *N*_+_, the surface density of receptors *σ_R_* and *χ*, a parameter related to the strength of the ligand-receptor bond. Furthermore, *z* is the so-called activity in solution^126^ of the multivalent construct, which for all practical cases of interest is given by *z* = *v*_0_*ρ_B_*, where *ρ_B_* is the molar concentration of the multivalent construct in solution and *v*_-_ the adsorption volume. Within the Langmuir description used here, *θ* = *N_ads_*/ *M* can be interpreted as the binding probability of a single multivalent particle binding to the surface^79^. In Eq 1, in contrast to the more general definition^79^, we assumed that each binding site contains exactly the same number of receptors.

In order to calculate *Q*, we assume that each of the *N_L_* ligands in the multivalent construct can independently form a bond with each of the *N_R_* = *σ_R_A* receptors, where A is the area of a binding site on the surface. In this case, the total bound partition function is given by:

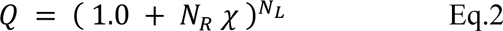

Where *χ* is related to the ligand-receptor dissociation constant *K_D_* by ^80^:

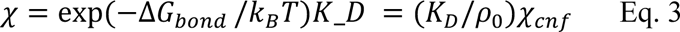

where Δ*G_bond_* is the effective bond energy, *ρ*_0_ = 1*M* is the standard molar concentration and 0 < *χ_cnf_* < 1 a (bond-weakening) correction that arises from the fact that the ligands are not free in solution, but attached to a scaffold (i.e. dextran) that limits their translational and rotational freedom. Although the exact value of *χ_cnf_* depends on details of how the ligands are grafted to the particle^80^, in our experiments it is the same across all systems, since the mixture of ligand (IgG) and the scaffold (dextran) is kept consistent within the same experiment.

*N_ads_* as described by Eq.1 has the form of a Hill-equation with an exponent equal to *N_L_*. Qualitatively, on a logarithmic scale the adsorption curve thus resembles a smoothened step function, with *N_ads_* ≈ 0 until the receptor density reaches a certain critical value, 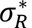, after which *N_ads_* increases very rapidly (at least until it reaches its maximum value, *M*). This is the behaviour we observed in our experiments (Figure 5C, Figure 6A).

To estimate the value of the bond dissociation constant (*K_D_*), we note that in the partition function *Q*, and thus also in the expression for the number of adsorbed particles *N_ads_*, *N_R_* (or *σ_R_*) and *χ* only appear as a product, not alone. Although the expression for *Q* is derived assuming independent ligands, more general treatments that do not make these assumptions^81,83^ still lead to expressions where only the *N_R_χ* product appears, making the following conclusions quite robust. Because these two quantities only appear as a product, mathematically, this means that when plotting log *N_ads_* vs log *σ_R_*, multiplying *K_D_* by an arbitrary (positive) factor *γ*, simply rigidly shifts all points of the curve by exactly −log *γ*. This happens for every single point on the curve, and thus also for the critical receptor density 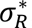 at which bound construct (IgG-dextran) begins to rapidly rise. In other words, if we plot *N_ads_* vs *σ_R_* for two systems *A* and *B* that differ only in their ligand-receptor pairs, in our case, different IgM antibody sequences, and we compare two points for these systems with the same value of *N_ads_*, we have:

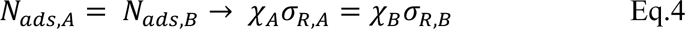

Or, using Eq.3 to connect with *K_D_* and rearranging the terms:

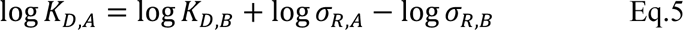

Notably, in Eq.5 the dependence on the configurational contribution *χ_cnf_* to the bond strength, which would be otherwise difficult to estimate, drops out because it is the same for two systems that only differ by the ligand-receptor pair used.

Thus, if the value of *K_D_* for a reference system *A* is known, Eq.5 allows us to calculate the *K_D_* value of any other ligand-receptor pairs by simply comparing the shift in the adsorption (i.e. IgM fluorescence threshold at which IgG-dextran binding occurs) on a logarithmic scale.

---

**Figure S1.**
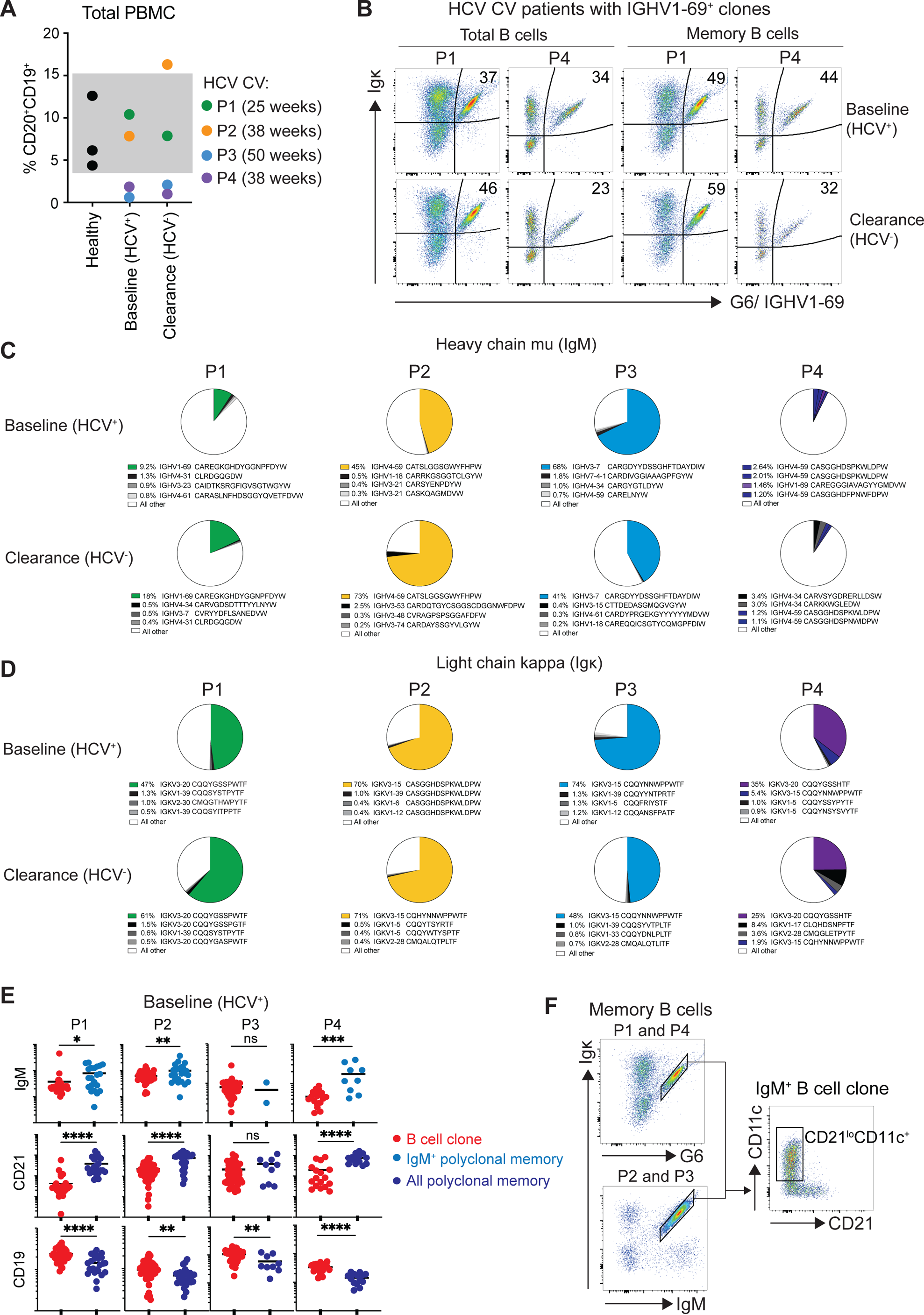
IgM^+^ B cell clonal expansions persist in the blood of cryoglobulinemic vasculitis patients following clearance of HCV. A. Frequency of B cells in total live PBMCs for 3 age-matched healthy controls and each patient at the baseline (HCV^+^) and clearance (HCV^−^) timepoints. Grey shading indicates the normal range in healthy individuals. B. Staining with the anti-idiotypic monoclonal antibody G6, showing the frequency of kappa-restricted IGHV1-69^+^ B cell clones in patients P1 and P4 at baseline (HCV^+^) and clearance (HCV^−^) timepoints. C. Total RNA was prepared from PBMCs, converted to cDNA, amplified with forward primers for the leader segments of all heavy V regions and a reverse primer for the mu constant region and amplified products analysed by Illumina MiSeq sequencing. The proportion of reads with the indicated V element and CDR3 amino acid sequence is shown for each patient and timepoint. The absence of the IGHV1-69 clonal expansion from the bulk sequencing of the patient P4 clearance timepoint was likely due to technical failure as the presence of the expanded IGHV3-20 light chain (D) and positive G6^+^/IGHV1-69^+^ staining by flow cytometry at both timepoints (B), indicated the same IGHV1-69/IGKV3-20 clone was present in the blood of patient P4 at clearance. D. Same as for C but using forward primers for all kappa V regions and a reverse primer for the kappa constant region. E. Results of indexed single cell flow staining and RNA sequencing at baseline in each patient, comparing mean fluorescence intensity of IgM, CD21 or CD19 on single cells (denoted by dots) identified as belonging to the B cell clone or polyclonal memory B cells based on heavy and light chain CDR3 immunoglobulin mRNA sequences. Unpaired Student’s t-test, ns; p ≥ 0.05 (not significant), * p < 0.05, ** p < 0.01, *** p < 0.001, **** p < 0.0001. F. Gating strategy to enumerate CD21^lo^ CD11c^+^ cells within the IgM B cell clone of each patient.

**Figure S2.**
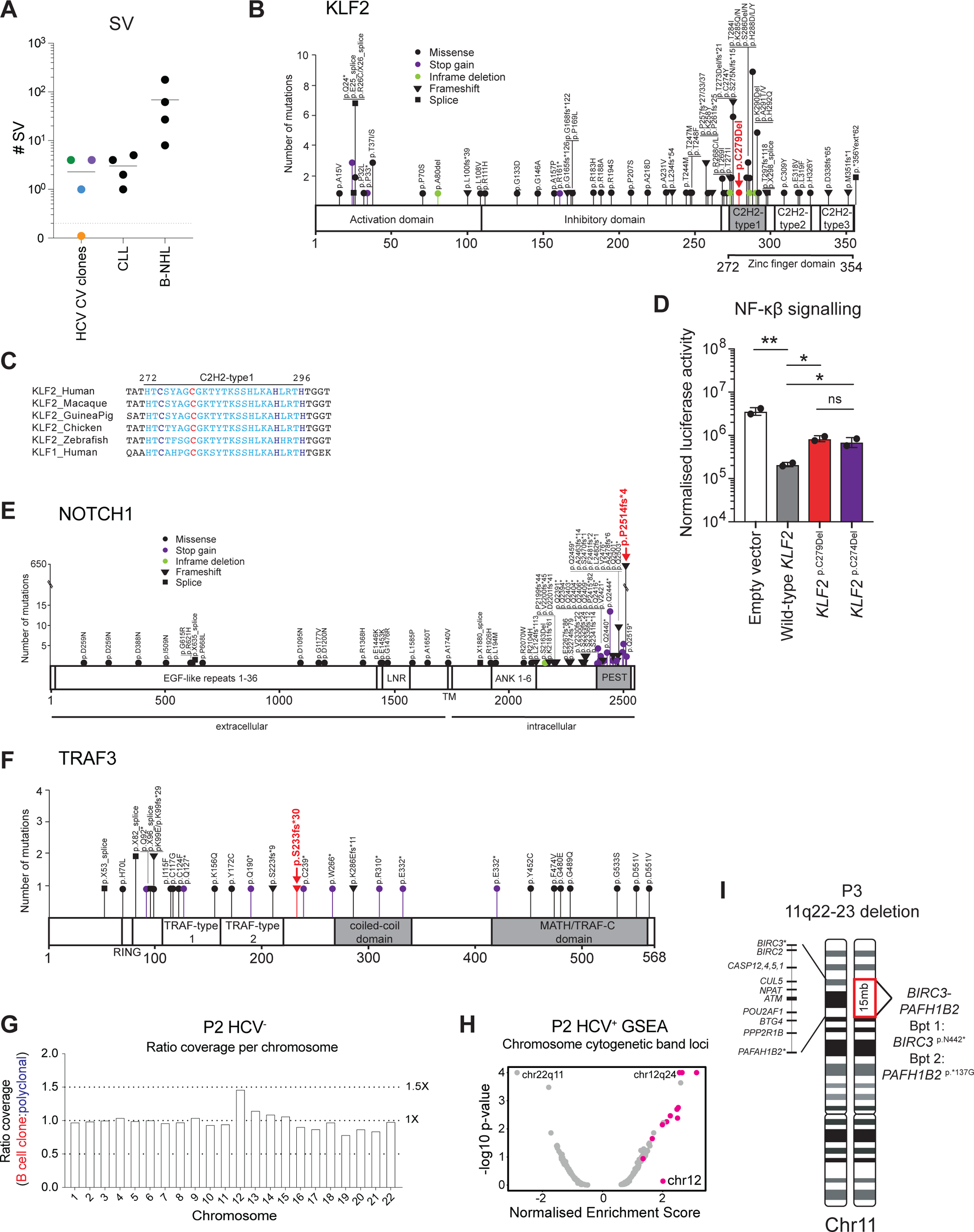
Lymphoma-associated somatic mutations in the IgM^+^ clone of each patient. A. Total number of chromosome structural variants (SV) including SV at immunoglobulin (IG) loci detected in each HCV-CV patient’s clone (P1-P4) compared to 4 representative IGHV-mutated CLL samples and 4 B cell non-Hodgkin’s lymphomas. Related to Figure 2F. B. *KLF2* somatic mutations previously reported in B cell lymphomas. The *KLF2^p.C^*^279^*^Del^* mutation identified in the IGHV1-69 clone of patient P1 is indicated in red. C. Amino acid alignment of KLF2 first zinc finger domain across species and with the paralogous human KLF1 zinc finger. Light blue indicates residues of the first C2H2 domain, red indicates the deleted cysteine (C) (position 279) and dark blue indicates the other cysteine (C) (position 274) and two histidine (H) residues (positions 292 and 296) that coordinate zinc ion binding. D. Luciferase reporter assay to determine the impact of KLF2 variants on NF-κB signalling. HEK293T cells were co-transfected with plasmid DNA encoding NF-κB luciferase, control reporter plasmid (β-galactosidase) and either empty plasmid (empty vector; white), wild-type KLF2 (grey) or KLF2 variant (*KLF2*^p.C279Del^; red, *KLF2*^p.C274Del^; purple). Transfected HEK293T cells were pulsed with TNFα two hours before harvest. NF-κB luminescence was normalised against β-galactosidase luminescence to control for transfection efficiency. Samples were run in duplicate. Unpaired student’s t-test, ns; not significant, * p<0.05, ** p<0.01. E. *NOTCH1* somatic mutations previously reported in B cell lymphomas. The *NOTCH1^p.P2514fs*^*^4^ mutation identified in the IGHV1-69 clone of patient P4 is indicated in red. F. *TRAF3* somatic mutations previously reported in B cell lymphomas. The *TRAF3^p.S233fs*^*^30^ mutation identified in the IGHV1-69 clone of patient P4 is indicated in red. G. Ratio of whole genome sequencing (WGS) average read depth per chromosome (chr 1-22) in patient P2’s clone and polyclonal memory B cells from the clearance timepoint (HCV^−^). H. Normalised gene set enrichment analysis (GSEA) score for chromosome location (cytogenic bands) of differentially expressed mRNAs in single cell RNA-seq comparing patient P2’s clone (*n*=110 cells) and polyclonal memory B cells (*n*=54 cells). Chromosome 12 cytogenetic bands shown in pink. I. Schematic of the 11q22-23 deletion in the IGHV3-7 clone patient P3 identifying the breakpoints in *BIRC3* and *PAFH1B2* and key protein coding genes located in the deleted segment on chromosome 11 (chr 11).

**Figure S3.**
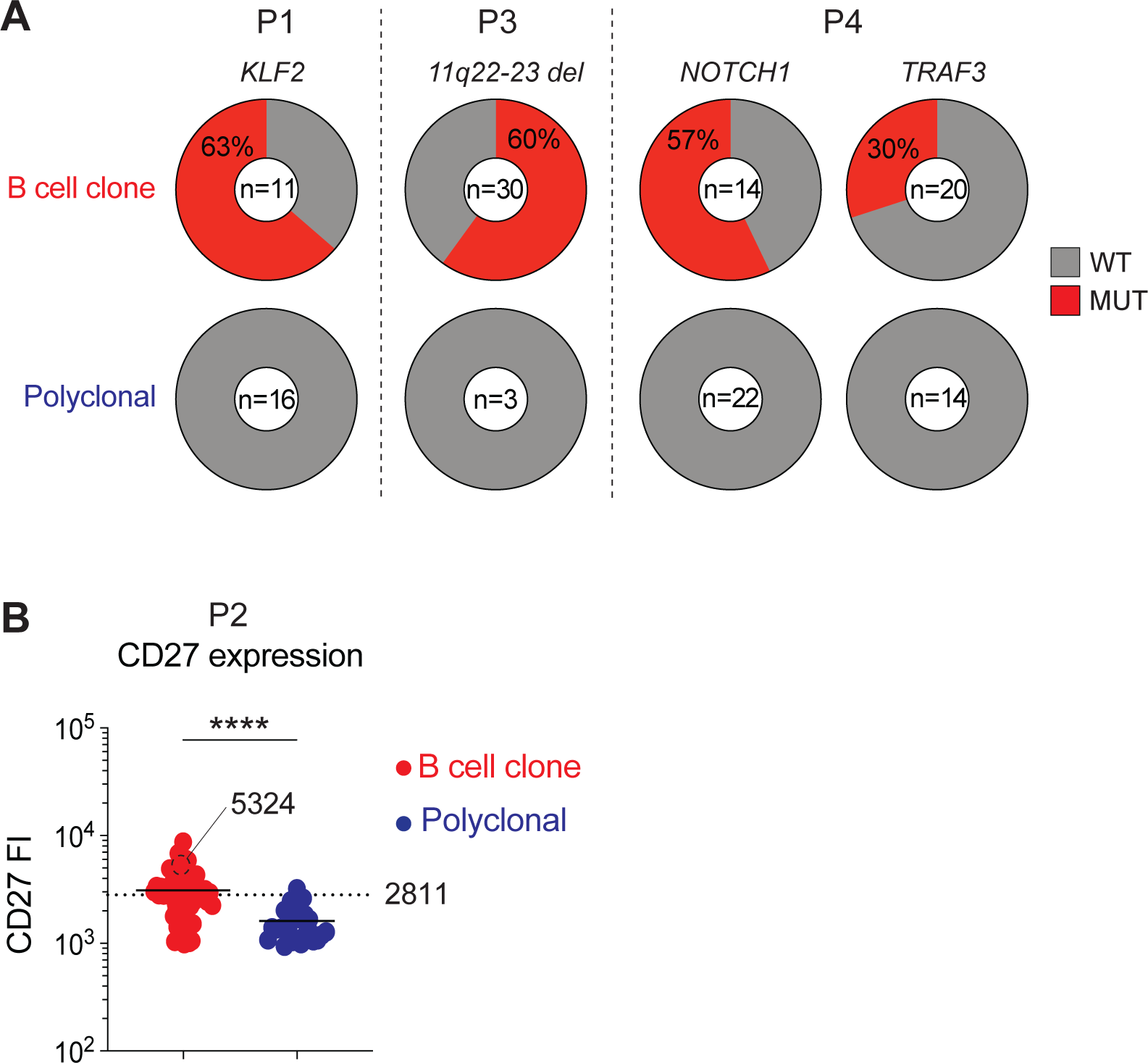
The somatic mutations are confirmed in single clonal B cells. A. Confirmation of the somatic mutations in individual clonal cells for patients P1, P3 and P4 by PCR and Sanger sequencing. For patients P1 and P4, the number of single cells with a high confidence mutant or wild-type allele call are indicated in the centre of the piegraph. For patient P3, the number of single cells positive for a PCR product for the control gene (*SEC23IP*) and/or 11q22-23 deletion are indicated at the centre of the piegraph. MUT = mutant allele, WT = wildtype allele. B. Indexed single cell flow cytometric data showing CD27 fluorescence intensity (FI) on individual clonal B cells (red) or individual polyclonal memory B cells (dark blue) from patient P2. B cell clonality was annotated based on heavy and light chain CDR3 immunoglobulin mRNA sequences. Black dotted line indicates two standard deviations above the mean of polyclonal B cells at the FI threshold of 2811 used identify cells with high CD27 expression (CD27^high^) in Figure 3E. Clonal single cell with black outline identifies the single cell from P2’s clonal tree lacking the IGHV4-59 S92T mutation with (CD27 FI 5324), as shown in Figure 3E.

**Figure S4.**
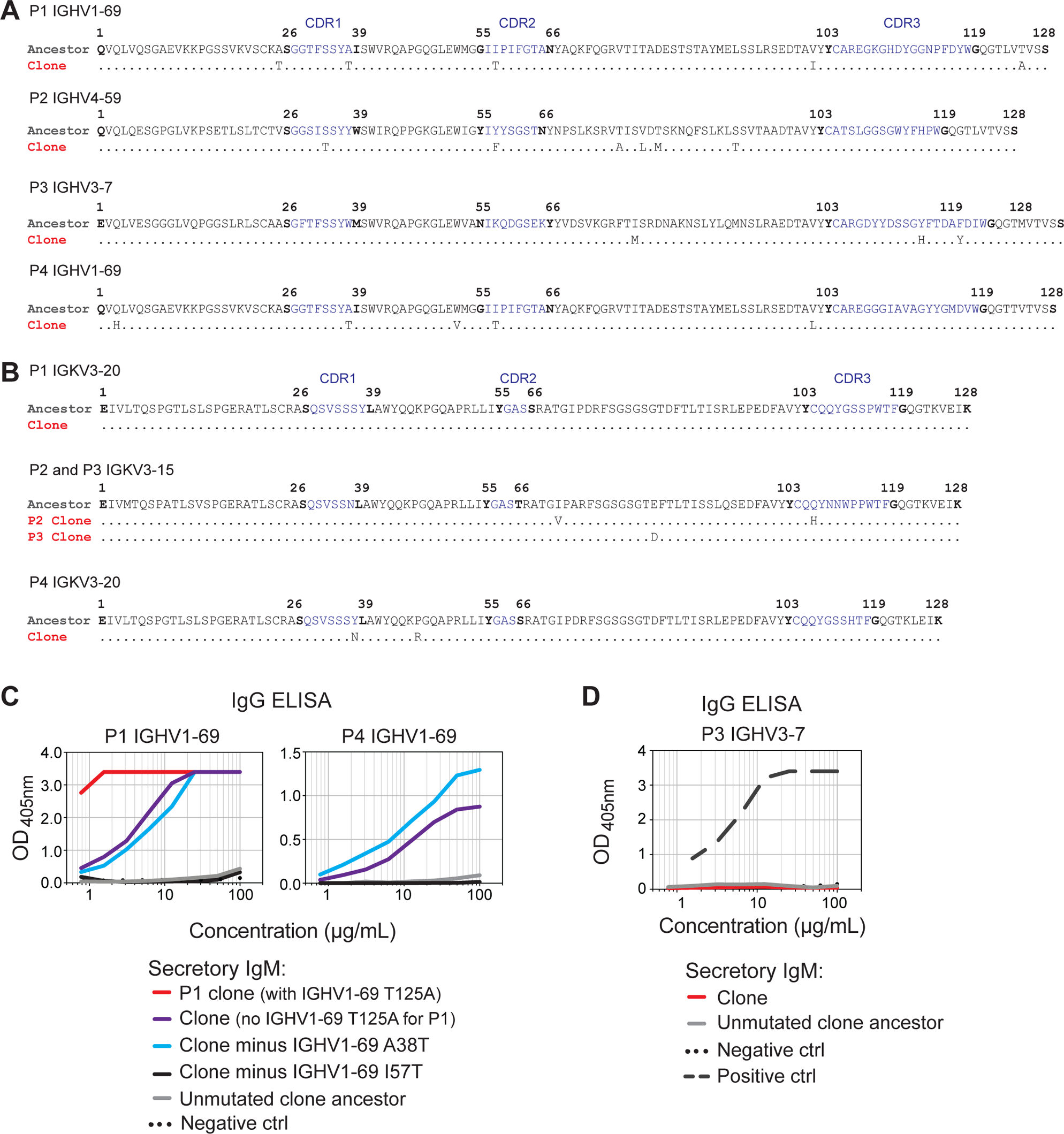
Self-reactivity of the mutated and unmutated ancestor clonal IgM immunoglobulin sequences. A. Amino acid sequence alignment of predominant clonal heavy chain *V(D)J* sequence in each patient (“clone”) and the inferred unmutated ancestor (“ancestor”). IMGT numbering, CDRs in blue with flanking residues bold. Period (.) indicates identity. B. Alignment of the predominant clonal (“clone”) and inferred unmutated (“ancestor”) light chain *VJ* sequences. IMGT numbering, CDRs in blue with flanking residues bold. Period (.) indicates identity. C. IgG binding in ELISA by synthesised secretory IgM corresponding to the predominant clonal sequence in patient P1 with (red) and without the T125A mutation (purple), predominant clonal sequence in P4 (purple), with all mutations reverted to ancestral sequence (grey), or the A38T mutation (light blue) or I57T mutation (black) selectively reverted. Negative control (black dotted) was anti-HCV broadly neutralising antibody AR3C expressed as secretory IgM. Representative of 2 independent experiments, run in duplicate. D. Synthesized secretory IgM corresponding to the predominant clonal sequence (red) of patient P3 and unmutated ancestor (grey) tested at the indicated concentrations for binding to immobilised human IgG by ELISA. Negative control (black) was anti-HCV broadly neutralising antibody AR3C expressed as secretory IgM. Positive control (black dashed) is the clonal sequence of patient P4. Representative of 4 independent experiments, run in duplicate.

**Figure S5.**
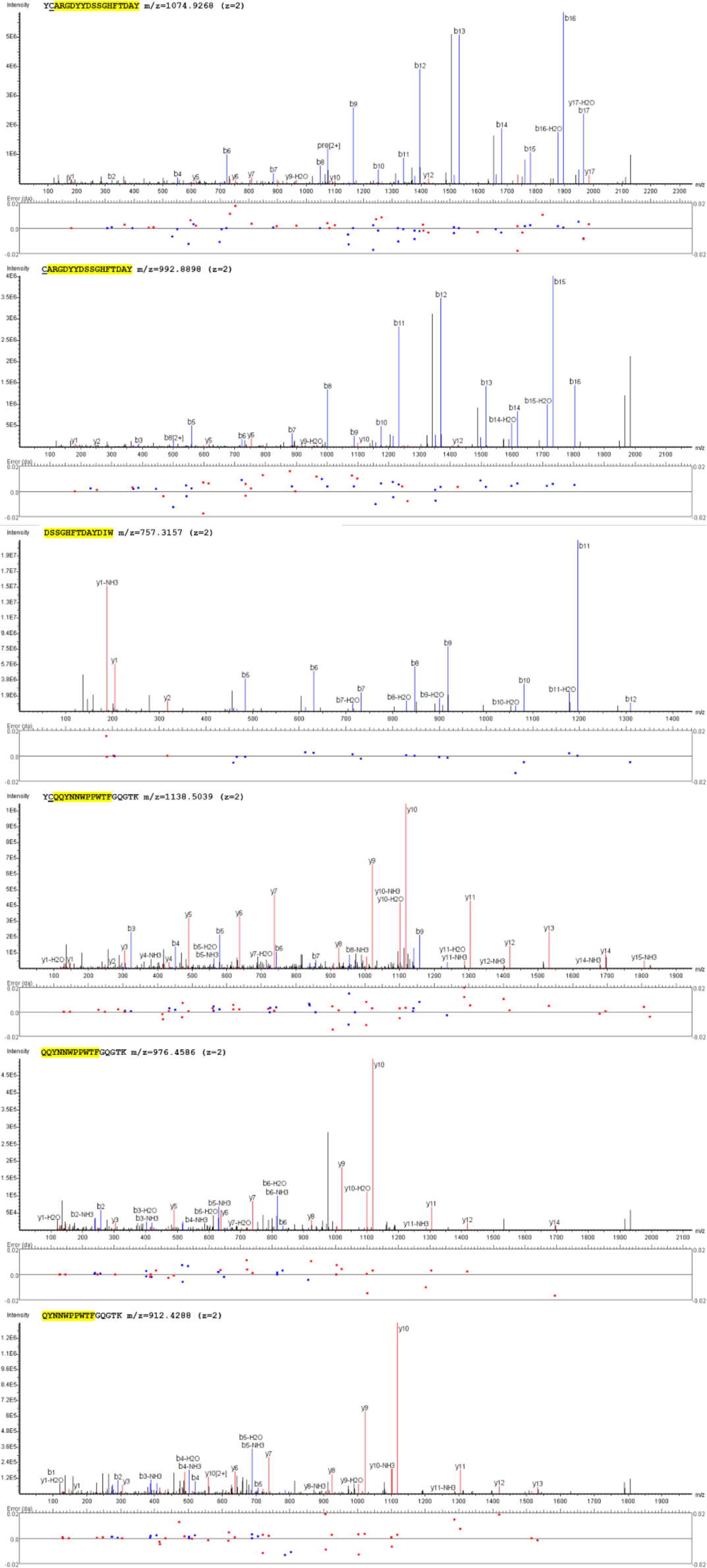
Representative annotated MS/MS spectra determining the amino acid sequence of HCDR3 and LCDR3 from the plasma RhF IgM antibody of patient P3. A. IgM-RF was immunoprecipitated from patient P3 plasma, heavy and light chains purified and cleaved into peptides sequenced by tandem mass spectrometry. The HCDR3 and LCDR3-containing peptides identified are highlighted in yellow. Underlined amino acid (C) indicates a carbamidomethylated cysteine. The sequences, m/z and z of each individual peptides are shown on the top of their annotated MS/MS spectra, and fragment ion error tolerance on the bottom of spectra. Matched b ions are indicated in blue and y ions are in red. m=mass, z=charge.

**Figure S6.**
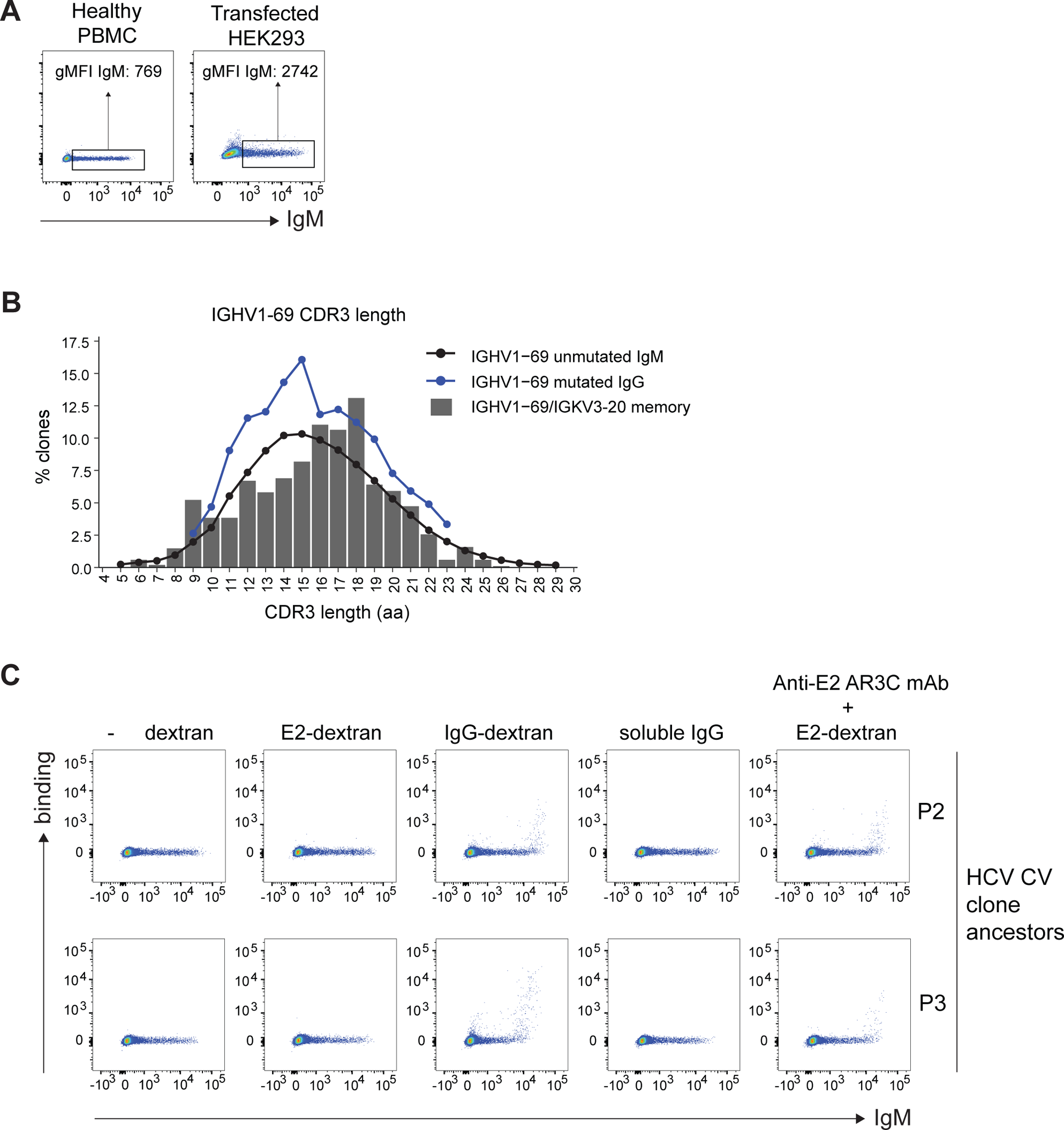
The RhF clonal ancestors bind multimerised IgG but not HCV E2. A. Comparison of the range of surface IgM expression on transfected cells and on PBMCs of a healthy donor stained and analysed in parallel. The IgM geometric mean fluorescence intensity (gMFI) for all IgM^+^ transfected cells (gate shown). Because the diameter of HEK293 cells is ∼2 times larger than the diameter of lymphocytes, HEK293 cells have ∼4 times greater surface area. The 3.6 fold higher IgM gMFI for the transfected HEK293 cells compared to human B cells is consistent with comparable concentrations of IgM per unit area of plasma membrane. B. Distribution of heavy chain IGHV1-69 CDR3 amino acid (aa) lengths for antibodies encoded by IGHV1-69/IGKV3-20 antibodies from 3 healthy donors (grey bars). The black and dark blue line denote the mean CDR3 amino acid (aa) length for IGHV1-69 from bulk repertoires from 61 healthy donors for unmutated IgM (naïve) and mutated IgG (memory) antibody sequences. C. Binding of empty dextran (first column), HCV E2 H77-dextran (second column), IgG-dextran (third column), soluble IgG (fourth column) and HCV E2 H77-dextran coated with anti-E2 AR3C monoclonal IgG1 antibody, to cells expressing the unmutated ancestors of patient B cell clone antibodies (P2,P3) as membrane IgM. HEK293 cells are transiently transfected and stained in parallel.

**Text from Figure S7.**

B. From Methods, we have the following equation linking membrane IgM threshold of binding (*σ_R_*) and affinity constant (*K*_D_).

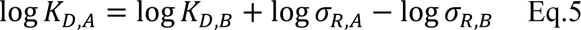

C. Using Eq.5 and membrane IgM IgG binding threshold fluorescence intensity as shown in Figure 6B, and using TJ4 with known IgG affinity constant (*K*_D,A_ = 11.2 x 10^−4^ M) to calculate the estimated affinity constant (*K*_D_) for the HCV B cell clone of patient P1:

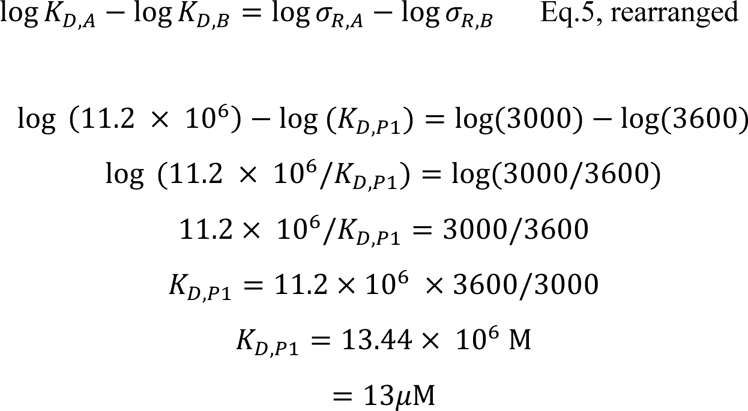

**Figure S7.**
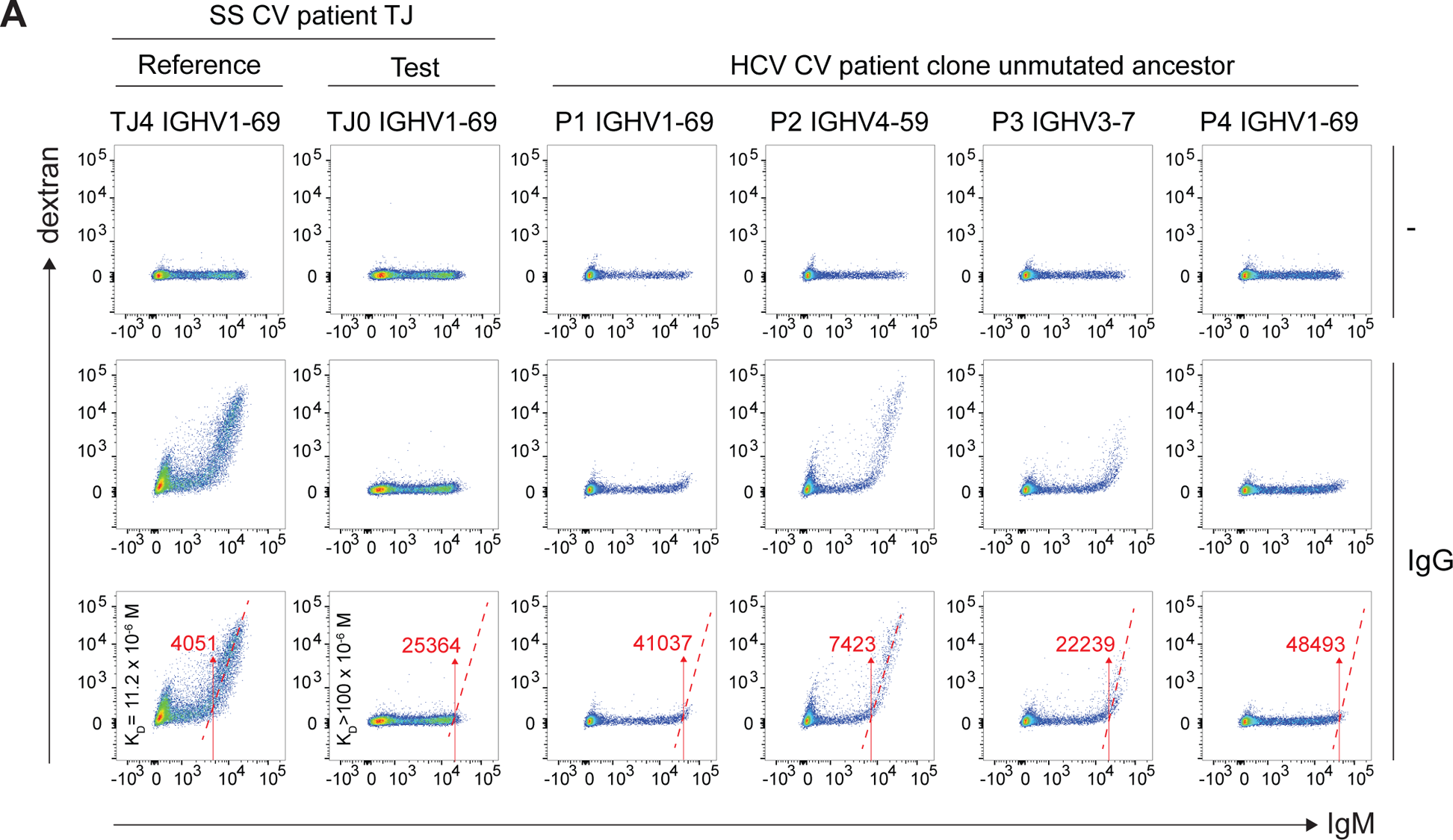
Equation linking membrane IgM binding threshold to affinity constant (K_D_) A. Binding of empty dextran (top row) and IgG-dextran (bottom two rows) to cells expressing a IGHV1-69/IGKV3-20 IgM RhF cryoglobulin from a Sjogrens’s syndrome patient with vasculitis (SS CV) (TJ4) and it’s unmutated ancestor (TJ0) with known affinities to IgG (as indicated), alongside B cell clone unmutated ancestor antibodies from patients P1-P4 as membrane IgM. HEK293 cells are transiently transfected and stained in parallel. The fluorescence intensity for membrane IgM at which IgG-dextran binding occurs is shown by the number and arrow in red (bottom row only). B. Equation describing the relationship between membrane IgM threshold of binding and affinity constant (K_D_), related to Figure 6C and Methods (Eq 5). C. Example calculation using the equation shown in A (Eq 5) to estimate the affinity (K_D_) of the mutated RhF of P1 using TJ4 as the reference (TJ4, K_D_ = 11.2 x 10^−6^ M, IgM fluorescence intensity = 3000) and the IgM fluorescence intensity threshold for IgG-dextran binding for P1 (P1 IGHV1-69, IgM fluorescence intensity = 3600). IgM fluorescence intensities as shown in Figure 6B.

